# Cancer-associated fibroblasts actively compress cancer cells and modulate mechanotransduction

**DOI:** 10.1101/2021.04.05.438443

**Authors:** Jorge Barbazan, Carlos Pérez-González, Manuel Gómez-González, Mathieu Dedenon, Sophie Richon, Ernest Latorre, Marco Serra, Pascale Mariani, Stéphanie Descroix, Pierre Sens, Xavier Trepat, Danijela Matic Vignjevic

**Author notes:** These authors contributed equally to this work (Jorge Barbazan, Carlos Pérez-González).

## Abstract

During tumor progression, cancer-associated fibroblasts (CAFs) accumulate in tumors and produce excessive extracellular matrix (ECM), forming a capsule that enwraps cancer cells. This capsule is a barrier that restricts tumor growth leading to the buildup of intratumoral pressure. Combining genetic and physical manipulations *in vivo* with microfabrication and force measurements *in vitro*, we found that the CAFs capsule is not a passive barrier but instead actively compresses cancer cells using actomyosin contractility. Cancer cells mechanosense CAF compression, resulting in an altered localization of the transcriptional regulator YAP. Abrogation of CAFs contractility *in vivo* leads to the dissipation of compressive forces and impairment of capsule formation. By mapping CAF force patterns in 3D, we show that compression is a CAF-intrinsic property independent of cancer cell growth. Supracellular coordination of CAFs is achieved through fibronectin cables that serve as scaffolds allowing force transmission. Our study unveils that the contractile capsule actively compresses cancer cells, modulates their mechanical signaling, and reorganizes tumor morphology.

---

Cancer progression is the result of complex interactions between cancer cells and their microenvironment^1^. Cancer cells continuously sense signals from the surroundings that could be either biochemical, such as soluble molecules or membrane receptors on stromal cells, or physical, including stiffness, the microarchitecture of the surrounding extracellular matrix (ECM), fluid pressure and solid stress^2^. Solid stress is a consequence of the proliferation of cancer cells against a viscoelastic boundary, the stroma, which resists tumor growth and prevents its expansion, resulting in the buildup of internal pressure^3,4^.

One of the most abundant cell types in the stroma are cancer-associated fibroblasts (CAFs)^5^. CAFs play multiple roles in cancer progression, promoting cancer cell survival and proliferation and modulating cancer invasion and immune response^5,6^. CAFs are highly contractile cells that serve as the main producers of the ECM^7,8^. Together with the ECM, they form a capsule around the tumor^9,10^. Is this capsule just a barrier that passively opposes tumor growth as generally assumed, or does it have an active role in the generation of tumor stresses? Here we investigated the role of CAFs’ contractility in capsule formation and how CAFs mechanically interact with cancer cells.

To study the organization of CAFs in tumors, we used a transgenic mouse model that spontaneously develops tumors in the intestinal epithelium due to the expression of an activated Notch1 (NICD) and the deletion of the tumor suppressor p53^10^ (hereafter referred to as N/p53/mTmG) (**Extended Data Fig. 1a**). Cancer cells were visualized through the expression of a nucleus- and membrane-targeted GFP, while all other cells expressed a membrane-targeted tdTomato. The stroma, composed of CAFs and extracellular matrix (ECM), formed a thick capsule enveloping the tumor. Interestingly, the stroma penetrated the tumor, compartmentalizing it into smaller clusters (**Extended Data Fig. 1b**; **Supplementary Video 1**). Each cluster was enwrapped with aligned CAFs forming intratumoral capsules. This was remarkably similar to the typical organization of human colorectal cancers^9^. Given that intratumoral capsules were rich in phosphorylated, thus active, myosin II (**Fig. 1a**), we wondered whether CAFs contractility could play a role in tumor compartmentalization.

**Fig. 1.**
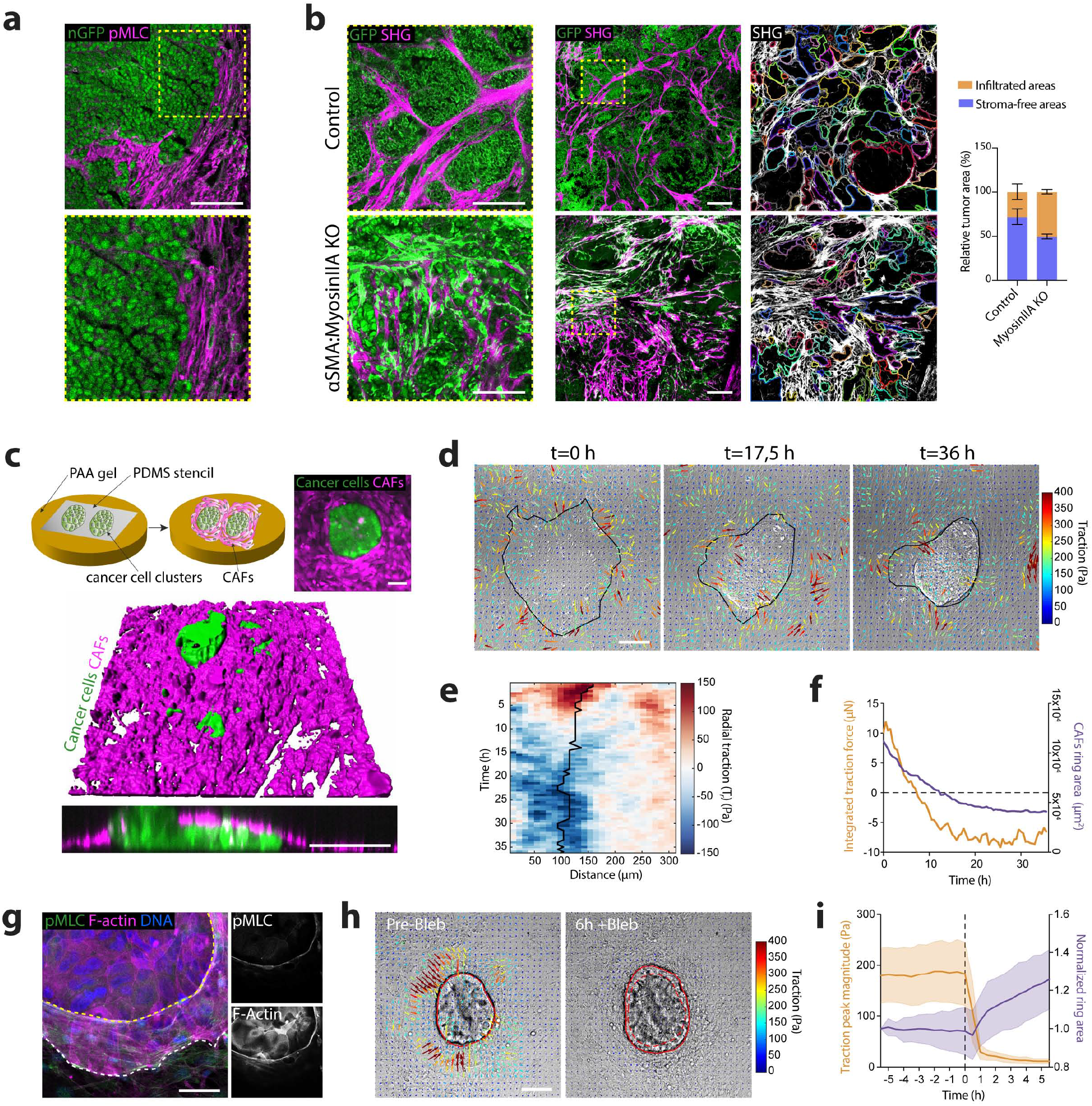
CAFs contractility is required for capsule formation and tumor compartmentalization. **(a)** Phospho-Myosin Light chain (pMLC) staining (magenta) in a tumor tissue section from N/p53/mTmG mice. Tumor cells express a nuclear GFP tag (nGFP). Scale bar 100µm. **(b)** Tumors generated after subcutaneous engraftment of N/p53 organoids into control mice and mice containing myosin IIA-knockout CAFs. Middle panel: Cancer cells (nuclear GFP), Myosin IIA KO CAFs (membrane-GFP, green). Collagen I revealed by SHG (magenta). Scale bar: 300µm. Left panel, magnified boxed regions. Scale bar, 100 µm. Right panel, automatic segmentation of stroma-free areas based on SHG signal. Right bar chart: quantification (%) of stroma-free (purple) or infiltrated (orange) tumor areas. Data represented as mean ± SD, N=2 mice per group, injected weekly with tamoxifen. **(c)** Top panel, schematic representation of in vitro co-culture protocol, and representative image of the resultant tissue organization (right panel): a cluster of primary cancer cells (Cell tracker, green) is surrounded by CAFs isolated from PDX (Cell tracker, magenta). Scale bar, 100 µm. Lower panel, 3D rendering of co-cultures after 48h in culture. Lower panel, orthogonal Z section. Scale bar: 100µm. (**d**) Representative traction maps overlaid on a DIC image of a cancer cell-CAFs co-culture, evolving over time. Black line represents the contour of the cancer cell cluster Scale bar, 100 µm. **(e)** Representative kymograph of circumferentially averaged radial tractions as a function of the distance to center of the cancer cell cluster (outwards pointing tractions are defined as positive, and inwards pointing tractions are negative). Solid black line represents the boundary of the cancer cell cluster. **(f)** Time evolution of the radial traction peak near the boundary of the cancer cell cluster in E (orange) and CAFs ring area (purple). **(g)** CAFs-cancer cells co-culture stained for E-cadherin (green), F-actin (phalloidin, magenta) and DNA (DAPI, blue). Insets, magnified boxed region showing staining for active myosin (pMLC) and F-actin. Scale bar, 100 µm. **(h)** Traction maps overlaid on a DIC image of cancer cell-CAFs after ≈40h co-culture (Pre-Bleb) and 6h after addition of blebbistatin (6h+Bleb). Solid red line represents the contour of CAFs ring; dashed line represents CAFs ring contour before blebbistatin addition. Scale bar, 100 µm. **(i)** Average radial traction peak magnitude near the cluster cell boundary (orange) and CAFs ring area (purple) normalized to the initial ring size, as a function of time. Dashed black line represents the moment of blebbistatin addition. Data represented as mean ± SD, n=30 clusters from N=3 independent experiments.

To test this, we transplanted N/p53 tumor organoids into transgenic mice in which we conditionally ablated myosin IIA in aSMA-expressing cells (SMA-CreERT2; R26mT/mG; myosin IIA^fl/fl^). In this mouse model, cancer cells expressed nuclear GFP. All stromal cells, including CAFs with wild-type myosin IIA, expressed a membrane-targeted tdTomato. In aSMA-expressing cells, tdTomato switched to GFP concomitantly with the induction of myosin IIA knockout (**Extended Data Fig. 2**). In control mice, as in spontaneously developed tumors, CAFs generated intratumoral capsules that compartmentalized and confined cancer cells (**Fig. 1b**). These capsules were largely absent from mice in which CAFs lacked myosin IIA and, instead, cancer cells and CAFs appeared mixed. To quantify this, we segmented large tumor areas using the collagen signal visualized by second harmonic generation (SHG) imaging as a proxy for tumor-stroma boundary. We found that control tumors contained larger stroma-free areas than tumors containing myosin IIA depleted CAFs (**Fig. 1b**). These data show that CAFs’ contractility is required for the formation of intratumoral capsules, and thus segregation of cancer cells into compartments. To study the mechanism by which CAFs confine cancer cells, we followed a bottom-up approach and developed an in vitro co-culture system that recapitulates tumor organization. For this, we fabricated flat 11 kPa polyacrylamide gels coated with collagen I and we generated circular clusters (150µm radius) of cancer cells isolated from patient-derived xenografts (PDX) surrounded by CAFs (**Fig. 1c, Extended Data Fig. 3**). As in vivo, CAFs aligned to each other and parallel to the cluster boundary, encapsulating cancer cells and restraining their spreading. Surprisingly, after 8h, CAFs assembled into a multicellular ring that slid on top of the cancer cells. As the ring advanced, it deformed clusters, ultimately inducing the multilayering of cancer cells and forming a three-dimensional bud (**Fig. 1c, Supplementary Video 2**). At this point, the ring ceased to advance, and the bud remained stable. CAFs also encapsulated and induced budding of N/53 tumor organoids (**Extended Data Fig. 3a-c, Supplementary Video 3**). Notably, the same sequence of events was observed ex vivo, where CAFs and cancer cells spontaneously exited from PDX tumor fragments (**Extended Data Fig. 4d, Supplementary Video 4**). This shows that CAFs can form capsule-like structures to confine cancer cells even in reductionist 2D in vitro environments. The fact that this capsule deforms and reshapes tumor cell clusters suggests that CAFs are not just a passive barrier against cell spreading but rather an active entity that exerts forces on cancer cells.

To understand the mechanics of cancer cell-CAF interaction, we quantified tissue forces using traction force microscopy (TFM) (**Fig. 1d, Extended Data Fig. 5, Supplementary Video 5**). Forces fluctuated across the tissue but systematically accumulated at the boundary between the CAFs and the cancer cell cluster. Given the radial symmetry of the system, we decomposed tractions into radial (Tr) and tangential (Tt) components, respective to the cluster boundary. We circumferentially averaged these tractions for each timepoint and plotted them as a function of the distance to the cluster center to build kymographs (**Fig. 1e, Extended Data Fig. 5b**). We found that average tangential forces, which likely correspond to migration of CAFs around cancer cell clusters, remained low and constant over time (**Extended Data Fig. 5b-c**). In contrast, radial tractions were maximal at the boundary between cancer cells and CAFs (**Fig. 1d-e, Extended Data Fig. 5c**). Initially, radial tractions were positive (pointing away from cancer cells), probably due to CAFs crawling towards the cluster (**Fig. 1d-f**). Later, when CAFs aligned parallel to the cluster boundary, radial tractions became negative (pointing towards cancer cells). These negative radial forces progressively increased as CAFs formed the supracellular ring, slid on top of the cancer cells and induced budding. Once the bud stabilized, traction forces plateaued (**Fig. 1f**).

The observed traction dynamics are consistent with CAFs forming a contractile ring that closes on top of cancer cells by a purse-string mechanism^11^. Ring closure builds up tension that, when transmitted to the substrate at the boundary with the cancer cell cluster, generates inward-pointing tractions (**Extended Data Fig. 5a**). Cluster deformation suggests that CAFs rings also transmit forces to the cancer cells. CAF closure applies a shear stress that induces cancer cells multilayering and bud formation. The integrated traction force plateaus prior to ring stabilization (**Fig 1f**), which indicates that a fraction of the ring tension is resisted by bud compression (**Theoretical Nore 1, Supplementary Figure 1**). At this point bud size remains stable over time. A simple elasto-plastic model, assuming compression-dependent cellular rearrangement shows that such buds may indeed be stable for a limited range of line tension (**Theoretical note 1**).

Similar to in vivo intratumoral capsules (**Fig. 1a**), in vitro CAF rings were enriched in phosphorylated myosin II (**Fig. 1g**). This suggests that ring constriction is driven by actomyosin contractility. Indeed, inhibition of contractility using either blebbistatin to inhibit myosin II (**Fig. 1h)** or Y27632 to inhibit ROCK (**Extended Data Fig. 5d**), induced a fast relaxation of CAF rings, almost complete disappearance of traction forces, and the flattening of the bud (**Fig. 1h-i, Extended Data Fig. 5d-e, Supplementary Video 6**). Similarly, myosin IIA knockout CAFs exerted significantly lower traction forces than control CAFs (**Extended Data Fig. 6a-c**). They failed to confine or deform cancer cell clusters even if forced to encircle them due to the experimental set up geometry (**Extended Data Fig. 6d-e**). Overall, our in vitro experiments suggest that CAFs intratumoral capsules not only passively confine, but also compress cancer cells.

To analyze the mechanical interactions between CAFs and cancer cells in tumors we performed two-photon laser ablations on ex vivo cultured thick tumor slices. First, we disconnected cancer cells from the surrounding stroma by performing ablations parallel to the edge of cancer cells (**Fig. 2a, Supplementary Video 7**). Upon ablation, we observed a rapid displacement of cancer cells towards the cut in tumor areas nearby the ablated region, whereas almost no displacement was observed in a distant control region (**Fig. 2b**). These data show that cancer cells are compressed in tumors. To address the tensional state of the CAFs, we also performed cuts perpendicular to the tumor boundary (**Fig. 2c**; **Supplementary Video 7**). Again, cancer cells displaced towards the cut, confirming that they were under compression, whereas CAFs recoiled away from the cut, indicating that they were under tension (**Fig. 2d**). Similar results were obtained using in vitro co-cultures of cancer cells and CAFs (**Fig. 2e-f; Supplementary Video 7**). Abrogation of contractility using blebbistatin or knocking out CAFs myosin IIA in tumor tissue slices decreased CAF recoil and directionality upon ablation (**Fig. 2f; Supplementary Video 8**). Even though there was a small displacement of cancer cells, directionality towards the cut was lost in tumors containing myosin IIA-knockout CAFs (**Fig. 2g, Supplementary Video 8**). As cancer cells had WT levels of myosin IIA, this shows that cancer cell compression is a direct consequence of CAFs contractility.

**Fig. 2.**
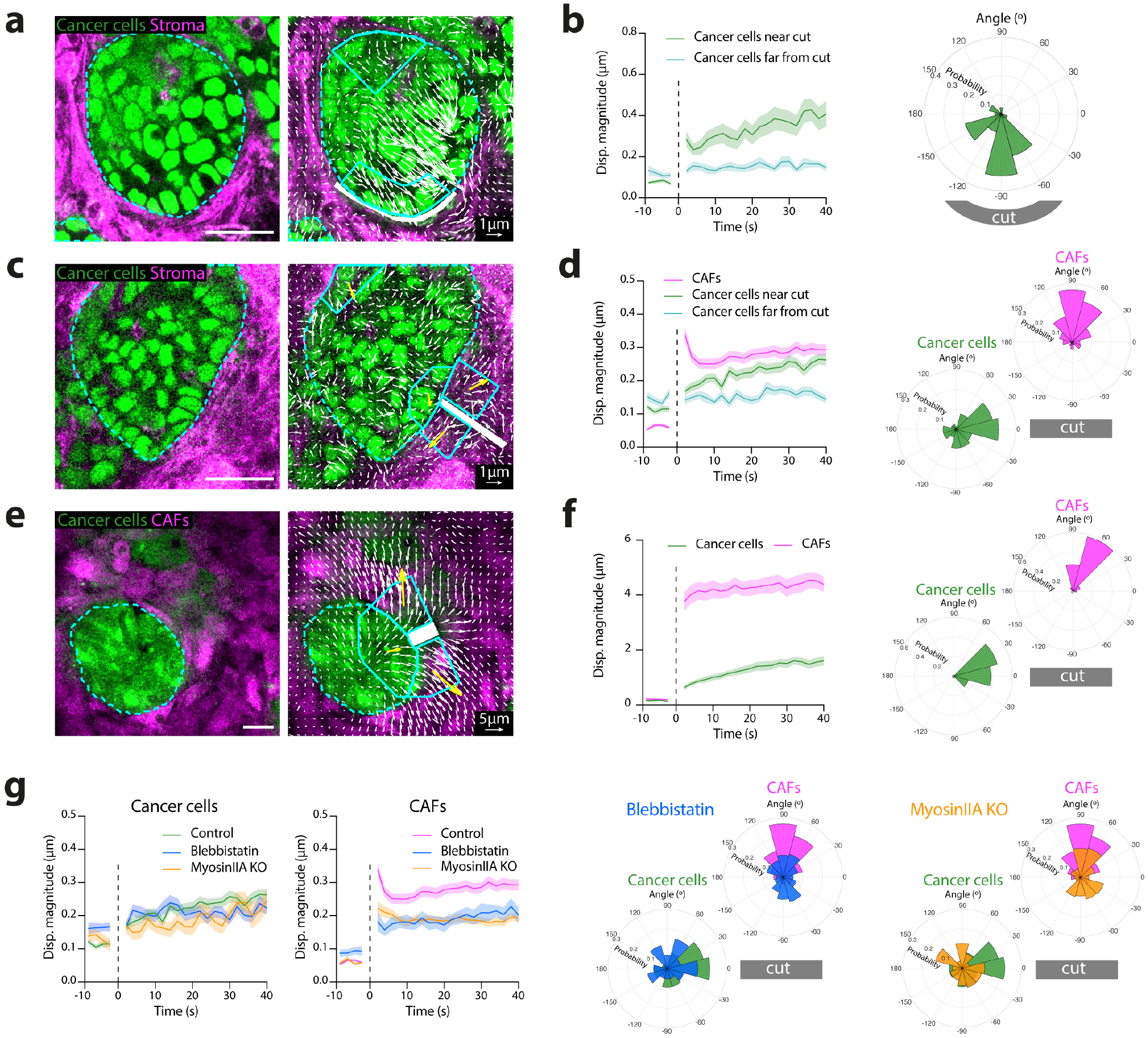
CAFs compress cancer cells using actomyosin contractility. **(a)** Montage showing images before (left) and after laser ablation (right) in fresh thick tumor slices. Cancer cells (nuclear-GFP, green), stroma (CAFs) (membrane-tdTomato, magenta). Dashed cyan line represents the contour of cancer cells. Solid cyan lines delineate 2 ROI, close and far away from the cut. Laser cut region is represented in white. White vectors, tissue displacement 40 seconds after ablation. Scale bar, 100 µm. Scale vector, 1 µm. **(b)** Displacement magnitude in 2 ROIs as a function of time after ablation (left). Vertical dashed line (t=0) indicates the ablation time. Polar histogram of displacement angle probability relative to the cut (right) 40 seconds after ablation. **(c)** Montage showing images before (left) and after laser ablation (right) in fresh thick tumor slices. Cancer cells (nuclear-GFP, green), stroma (CAFs) (membrane-tdTomato, magenta). Dashed cyan line represents the contour of cancer cells. Solid cyan lines delineate 4 ROI, close and far away from the cut in cancer cells and on both sides of the cut in CAFs. Laser cut region is represented in white. White vectors, tissue displacement maps 40 seconds after ablation. Yellow vectors, average displacement for each ROI (for visualization purposes, yellow vectors are not scaled). Scale bar, 100 µm. Scale vector, 1 µm. **(d)** Displacement magnitude of cancer cells and CAFs as a function of time after ablation (left). Vertical dashed line (t=0) indicates the ablation time. Polar histograms of displacement angle probability relative to the cut, for CAFs and cancer cells (right) 40 seconds after ablation. **(e)** Montage showing images before (left) and after laser ablation (right) in in vitro co-cultures. Cancer cells (Cell tracker, green), CAFs (Cell tracker, magenta). Dashed cyan line represents the contour of cancer cells. Solid cyan lines delineate 3 ROI, close from the cut in cancer cells and on both sides of the cut in CAFs. Laser cut region is represented in white. White vectors, tissue displacement 40 seconds after ablation. Yellow vectors, average cumulative displacement for each ROI (for visualization purposes, yellow vectors are not scaled). Scale bar, 100 µm. Scale vector, 5 µm. **(f)** Displacement magnitude of cancer cells and CAFs as a function of time (left) after ablation. Vertical dashed line (t=0) indicates the ablation time. Polar histograms of displacement angle probability relative to the cut, for CAFs and cancer cells (right) 40 seconds after ablation. **(g)** Displacement magnitude of cancer cells and CAFs as a function of time after ablation, in control tumor slices, tumor slices treated with blebbistatin or tumor slices contacting Myosin IIA KO CAFs (left). Vertical dashed line (t=0) indicates the ablation time. Polar histograms of displacement angle probability relative to the cut, for CAFs and cancer cells (right) 40s after ablation.

To address if cancer cells sense and functionally respond to CAF compression, we analyzed the cellular localization of a well-established mechanosensor, the transcriptional co-activator YAP, which shuttles in and out of the nucleus depending on the mechanical stress to which cells are subjected^12^. We measured YAP localization by quantifying the 3D correlation between DAPI and YAP signals (**Fig. 3**). A positive correlation indicates that YAP is preferentially in the nucleus, whereas a negative correlation reflects YAP cytoplasmic localization. We found that, in the absence of CAFs, YAP was mostly nuclear in cancer cells (**Fig. 3a**). Spatial confinement of cancer cells on micropatterns lead to an increase in cytoplasmic YAP. In the presence of CAFs, YAP localization was even more cytoplasmic. However, if CAFs lacked myosin IIA, YAP translocation to the cytoplasm was inhibited, demonstrating that CAF-mediated compressive forces are necessary to trigger YAP nuclear exit in cancer cells. Likewise, tumors with myosin IIA-knockout CAFs showed significantly increased nuclear YAP levels compared to control tumors (**Fig. 3b**). Altogether, these data show that CAFs, by compressing cancer cells, mechanically regulate YAP nuclear shuttling.

**Fig. 3.**
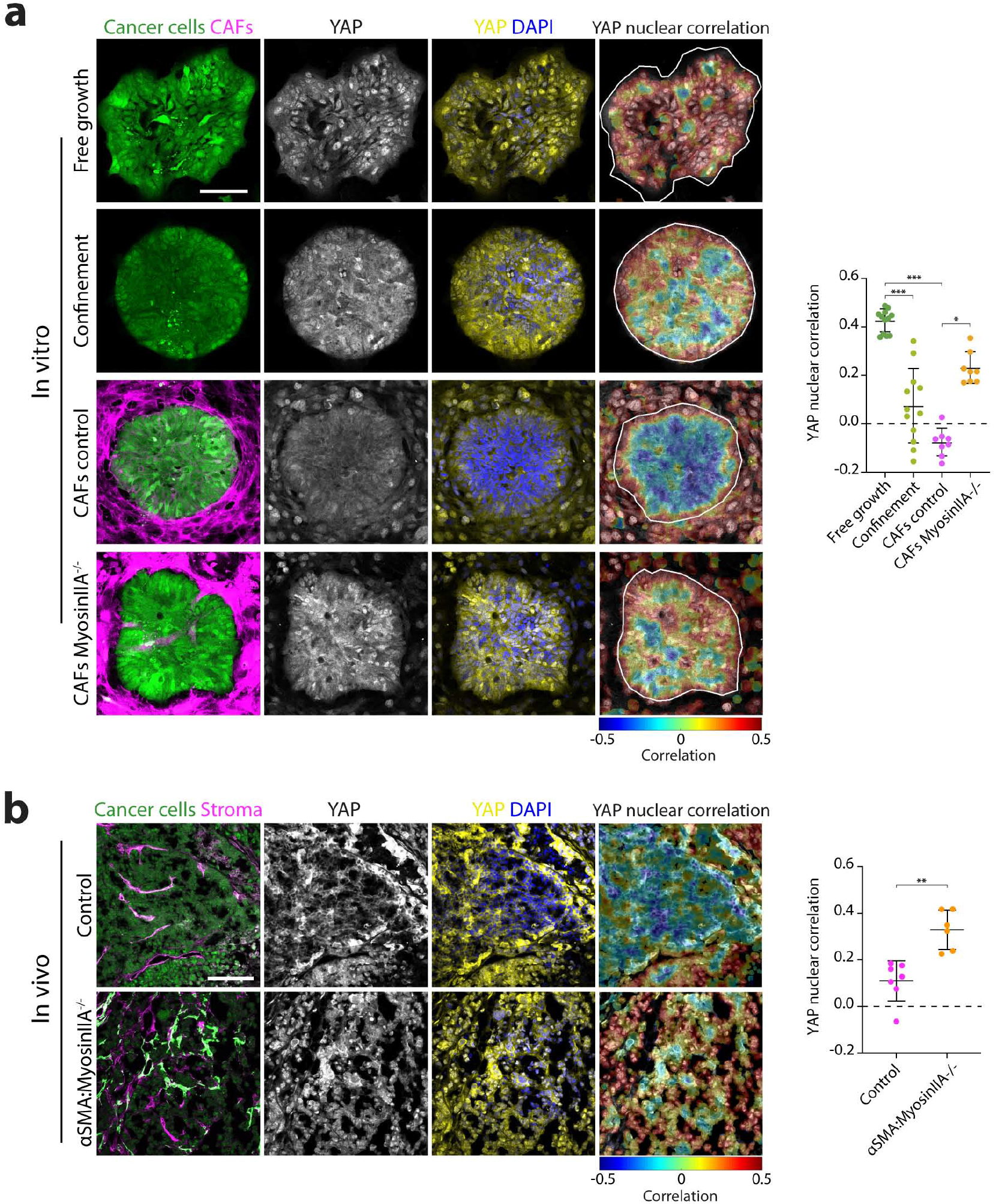
Cancer cells respond to CAFs compression through YAP nuclear translocation. (**a**) Cancer cells (Cell tracker, Green) cultured on an adherent substrate where they can freely spread, under confinement (PDMS stencils), or surrounded by control or Myosin IIA KO CAFs (F-Actin, phalloidin, magenta). Cells are stained for YAP (yellow) and DNA (DAPI, blue). Right panel, YAP nuclear correlation, ranging from 0.5 (red, high correlation) to -0.5 (blue, anti-correlation). White line outlines the boundary of the cancer cell cluster. Scale bar, 100 µm. Right dot plot, mean YAP nuclear correlation of cancer cells in vitro. Data represented as mean ± SD, n=8-12 clusters per condition, from N=3 independent experiments. * p<0.05, *** p<0.001, Kruskal-Wallis ANOVA test, Dunns multiple comparisons test. **(b)** Tumors from control or aSMA:Myosin IIA KO mice showing cancer cells (nuclear-GFP, green), CAFs expressing myosin IIA (membrane-tdTomato, magenta) and myosin IIA KO CAFs (membrane-GFP, green), stained for YAP (yellow) and DNA (DAPI, blue). Right panel, YAP nuclear correlation. Scale bar, 100 µm. Right dot plot, mean YAP nuclear correlation in cancer cells in vivo. Data represented as mean ± SD, each dot is the average of 3-6 image quantifications per tumor of n=7 control mice and n=6 myosin IIA KO mice, from N=2 independent experiments. ** p<0.1, Mann Whitney test.

We then asked whether the generation of compressive forces is a CAF intrinsic property or induced by cancer cells. To test this, we microfabricated 50µm radius soft polyacrylamide pillars (11kPa) to mimic the presence of cancer cells (**Fig. 4a, Extended Data Fig. 7a, Supplementary Video 9**). CAFs aligned parallel to the pillar border, assembled an actomyosin ring (F**ig. 4b-d**; **Supplementary Video 9**) and deformed the pillar (**Fig. 4e**). We measured pillar deformation using 3D particle image velocimetry and computed CAFs exerted forces using traction force microscopy^13^. This provided 3D traction force fields that we decomposed into a normal component (Tn, perpendicular to the pillar surface) and a tangential component (Tt, parallel to the pillar surface). Normal forces exhibited a larger magnitude than tangential forces (∼350 Pa vs ∼200 Pa on average) and were mostly compressive (negative) (**Fig. 4f-g, Extended Data Fig. 7b, Supplementary Video 10**). These forces decreased dramatically after inhibition of CAF actomyosin contractility using blebbistatin or Y27632 (**Fig. 4h, Extended Data Fig. 7c**). Altogether, these data show that CAFs can even compress inert materials and thus, compressive forces emerge from CAFs intrinsic contractility independently of the presence of cancer cells.

**Fig. 4.**
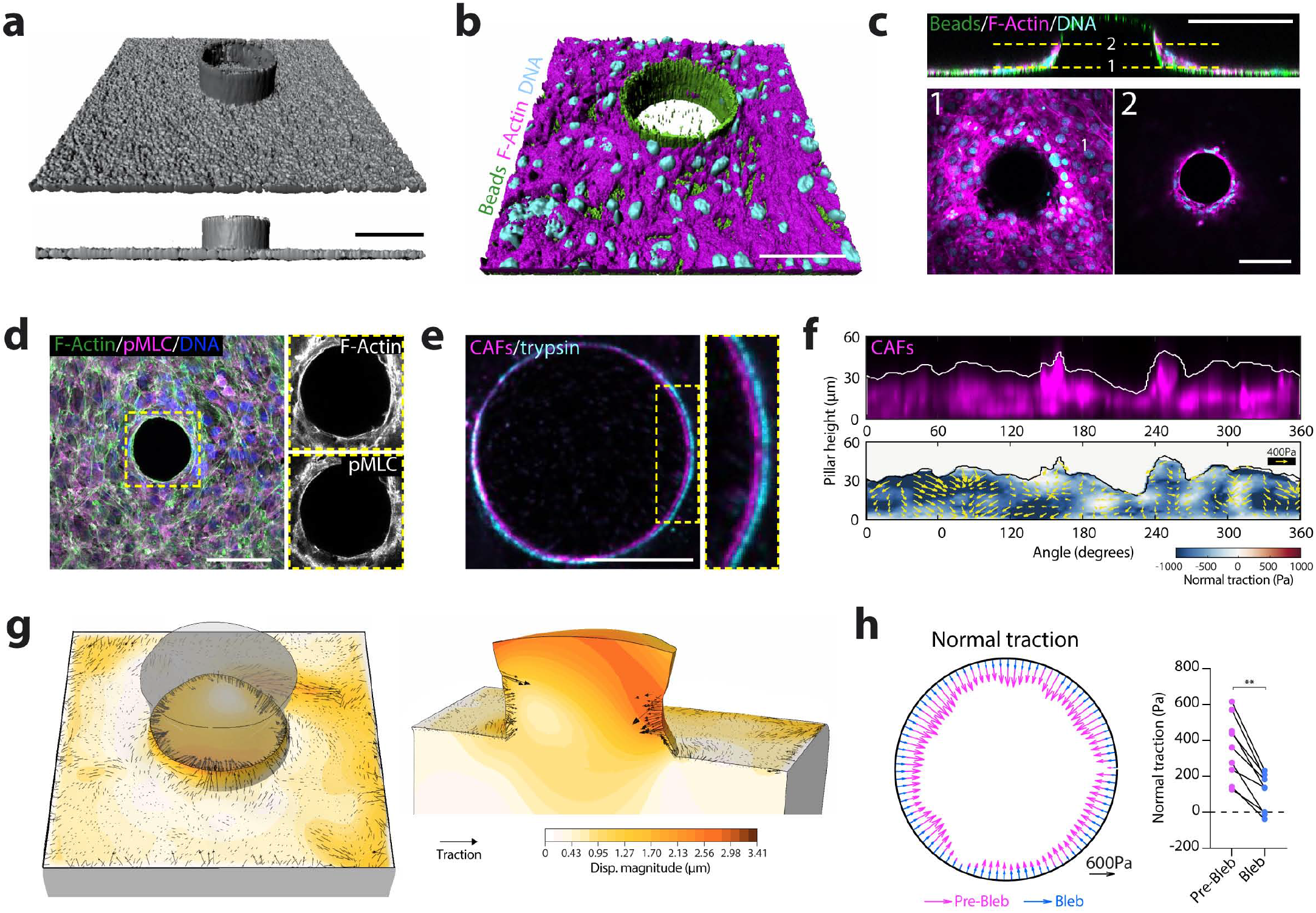
Compression is an intrinsic property of CAFs. **(a)** PAA pillars containing fluorescent beads (gray). Top, 3D view. Bottom, lateral view. Scale bar, 100 µm. **(b)** 3D rendering of CAFs stained for F-actin (phalloidin, magenta) and DNA (DAPI, cyan) surrounding a pillar (fluorescent beads, green). Scale bar, 100 µm. **(c)** Top, x-z pillar cross-section (fluorescent beads, green) surrounded by CAFs (F-actin, magenta; DNA, cyan). Dashed line represents two selected x-y planes shown at the bottom (1 and 2). Scale bars, 100 µm. **(d)** Maximum intensity projection of CAFs surrounding a pillar and stained for F-actin (phalloidin, green), active myosin (pMLC, magenta) and DNA (DAPI, blue). Inset, magnified boxed region showing staining for pMLC and F-actin. Scale bar, 100 µm. **(e)** Cross-section of a representative pillar at its base, in the presence of CAFs (magenta) and after removal of CAFs (trypsin, cyan). Inset, magnified boxed region. Scale bar, 50 µm. **(f)** Unwrapped pillar showing CAFs occupancy (top, magenta) and 3D traction force maps (bottom). Color map represents normal tractions (compression is defined as negative, pulling is defined as positive); yellow vectors represent tangential tractions. **(g)** 3D mapping of deformations (orange) and traction forces (black arrows) exerted by CAFs on a pillar. Top (left) and side (right) views. Deformations are magnified 5 times for visualization purposes. **(h)** CAFs normal tractions averaged across pillar height, on a representative pillar before (magenta) and after (blue) blebbistatin treatment. Scale vector, 600 Pa. Right dot plot, quantification of mean normal tractions for each pillar before and after blebbistatin. n=9 pillars, from N=2 independent experiments. ** p<0.1, Wilcoxon matched-pairs signed rank test.

To effectively compress and deform cancer cells, supracellular CAF rings should be sufficiently stable to maintain integrity over time. What mediates connections between CAFs in the ring? Mesenchymal cells, such as CAFs, generally lack stable cell-cell junctions. Instead, they are interconnected through N-cadherin zipper-like adhesions that allow cell-cell contacts while permitting fast neighbor exchange^14^. To test if N-cadherin mediates connections between CAFs in rings, we depleted N-cadherin using siRNAs (**Extended Data Fig. 8a**). Surprisingly, we found that N-cadherin-depleted CAFs were still able to compress pillars (**Fig. 5a**) and exerted the same levels of forces as control cells (**Extended Data Fig. 8b**).

**Fig. 5.**
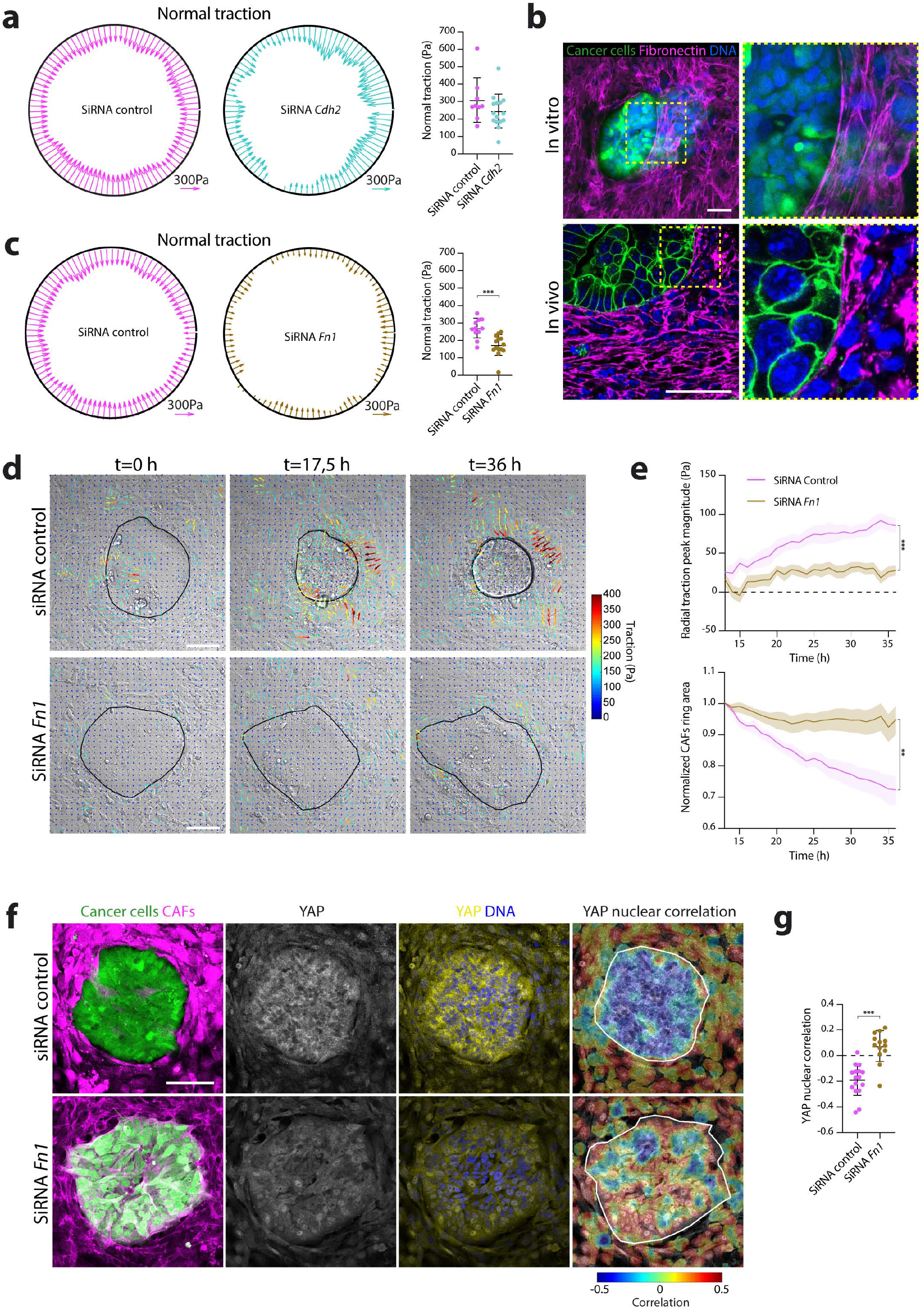
Fibronectin scaffolds allow CAFs supracellular coordination. **(a)** CAFs normal tractions averaged across pillar height, on a representative pillar for control (magenta) and N-cadherin depleted (cyan) CAFs. Scale vectors, 300 Pa. Right dot plot, quantification of mean normal traction per pillar. Data represented as mean ± SD. n=9 (control) and n=15 (N-cadherin knockdown) pillars, from N=3 independent experiments. Non-significant, Mann Whitney test. **(b)** Top left panel, cancer cells (cell tracker, green) and CAFs in vitro co-cultures, stained for fibronectin (magenta) and DNA (DAPI, blue). Right top panel, inset. Lower left panel, cancer cells (membrane GFP, green) and CAFs in tumors from N/ p53/mTmG mice, stained for fibronectin (magenta) and DNA (DAPI, blue). Scale bars, 50 µm. **(c)** CAFs normal tractions averaged across pillar height, on a representative pillar for control (magenta) and fibronectin depleted (brown) CAFs. Scale vectors, 300 Pa. Right dot plot, quantification of mean normal tractions per pillar. Data represented as mean ± SD. n=11 (control) and n=14 (Fibronectin knockdown) pillars, from N=3 independent experiments. *** p<0.001, Mann Whitney test. **(d)** Representative traction maps overlaid on a DIC image of cancer cell and CAFs control (top) or fibronectin depleted (bottom), evolving over time. Black solid line represents the contour of the CAFs ring. Scale bar, 100 µm. (**e**) Upper panel, radial traction peak magnitude at the boundary of the cluster for control (magenta) and fibronectin depleted (brown) CAFs, as a function of time. Lower panel, CAFs ring area normalized to the initial cluster size, as a function of time. Data represented as mean ± SEM. n=12 (control) and n=16 (Fibronectin knockdown) clusters, from N=3 independent experiments. *** p<0.001, ** p<0.1, Mann Whitney test for time point t=36. **(f)** Cancer cells (Cell tracker, Green) cocultured with control or fibronectin-depleted CAFs (F-Actin, phalloidin, magenta). Cells are stained for YAP (yellow) and DNA (DAPI, blue). Right panel, YAP nuclear correlation. White line outlines the boundary of the cancer cell cluster. Scale bar, 100 µm. **(g)** Dot plot, mean YAP nuclear correlation in cancer cells. Data represented as mean ± SD, n=16 (control) and n=13 (fibronectin knockdown) clusters, from N=3 independent experiments. *** p<0.001, Mann Whitney test.

In contrast to normal fibroblasts, CAFs produce excessive amounts of ECM proteins, especially fibronectin^9, 15-17^. Fibroblasts can use fibronectin to connect to each other using specialized cell-cell contacts, named stitch adhesions^18^. This led us to hypothesize an alternative mechanism for CAFs supracellular coordination indirectly via ECM. We observed that CAFs deposited abundant amounts of fibronectin both in vitro and in vivo (**Fig. 5b**). Consistent with the fact that CAFs contractility is required for fibronectin fibrillogenesis^15^, myosin IIA-knockout CAFs produced lower amounts of fibronectin (**Extended Data Fig. 8c-d**). The fibronectin network was isotropic and disorganized in the bulk of CAFs but became aligned at the boundary with cancer cells forming supracellular cables that spanned several CAFs (**Fig. 5b**). Similar reorganization of fibronectin network was previously seen in fibroblasts growing in macroscopically engineered clefts^19^. This led us to hypothesize that fibronectin could act as a mechanical scaffold for CAFs supracellular coordination and force generation during compression. Indeed, depletion of fibronectin in CAFs significantly reduced their ability to compress pillars (**Fig. 5c, Extended Data Fig. 8e**). Similarly, in the absence of fibronectin, CAFs could not stabilize multicellular rings, which led to a reduction in traction forces when co-cultured with cancer cells (**Fig. 5d-e, Supplementary Video 11**). Importantly, this loss of tractions is not due to impaired force generation because control and fibronectin depleted CAFs exerted similar tractions when cultured as single cells (**Extended Data Fig. 8f**). This shows that fibronectin is dispensable for force generation but required for intercellular force transmission. A direct consequence of this mechanical hindrance was the inability of CAFs to compress cancer cells (**Fig. 5d-e**) and thus induce the release of YAP from the nucleus (**Fig. 5f-g**). Altogether, these data show that fibronectin cables support force transmission between CAFs, allowing the mechanical coordination required to compress cancer cells.

Overall, we found that CAFs assemble intratumoral capsules stabilized by fibronectin scaffolds, allowing coordinated supracellular contraction. These contractile capsules actively compress cancer cells, triggering mechanical signaling and inducing reorganization of the tumor architecture. Thus, in contrast to the generally accepted concept, the tumor capsule is not just a passive barrier that prevents tumor expansion. Our data rather points towards capsules as active determinants of tumor solid stress. In light of previous observations, capsule-driven compression could favor tumor progression by promoting cancer cell stemness and proliferation^20^, as well as tumor budding, a poor prognosis factor for colorectal cancer patients^21-24^. These results support the use of targeted therapies against stromal contractility to improve the current efficacy of anti-cancer drugs.

## Acknowledgments

We thank all members of the DMV lab and G. Montagnac for helpful discussions and Raimon Sunyer for discussions on the microfabrication of PAA pillars. We acknowledge V. Fraisier and O. Renaud from the Cell and Tissue Imaging facility (PICT-IBiSA) and Animal Facility, Institut Curie.

## Author contributions

Conceptualization: JB, CPG, DMV

Methodology: JB, CPG, MGG, SR, EL, MS, PM

Data analysis: JB, CPG, MGG, EL, MD, PS

Funding acquisition: XT, DMV

Writing – original draft: JB, CPG, XT, DMV

Writing – review & editing: all authors

## Data and materials availability

Material and analysis tools are available upon request.

## References

1. J. A. Joyce, J. W. Pollard, Microenvironmental regulation of metastasis. Nat Rev Cancer 9, 239–252 (2009).

2. H. T. Nia, L. L. Munn, R. K. Jain, Physical traits of cancer. Science 370, (2020).

3. H. T. Nia et al., Solid stress and elastic energy as measures of tumour mechanopathology. Nat Biomed Eng 1, (2016).

4. T. Stylianopoulos et al., Causes, consequences, and remedies for growth-induced solid stress in murine and human tumors. Proc Natl Acad Sci U S A 109, 15101–15108 (2012).

5. E. Sahai et al., A framework for advancing our understanding of cancer-associated fibroblasts. Nat Rev Cancer 20, 174–186 (2020).

6. R. Kalluri, The biology and function of fibroblasts in cancer. Nat Rev Cancer 16, 582–598 (2016).

7. Y. Attieh, D. M. Vignjevic, The hallmarks of CAFs in cancer invasion. Eur J Cell Biol 95, 493–502 (2016).

8. J. Barbazan, D. Matic Vignjevic, Cancer associated fibroblasts: is the force the path to the dark side? Curr Opin Cell Biol 56, 71–79 (2019).

9. A. Glentis et al., Cancer-associated fibroblasts induce metalloprotease-independent cancer cell invasion of the basement membrane. Nat Commun 8, 924 (2017).

10. R. Staneva et al., Cancer cells in the tumor core exhibit spatially coordinated migration patterns. J Cell Sci 132, (2019).

11. S. R. Vedula et al., Mechanics of epithelial closure over non-adherent environments. Nat Commun 6, 6111 (2015).

12. T. Panciera, L. Azzolin, M. Cordenonsi, S. Piccolo, Mechanobiology of YAP and TAZ in physiology and disease. Nat Rev Mol Cell Biol 18, 758–770 (2017).

13. E. Latorre et al., Active superelasticity in three-dimensional epithelia of controlled shape. Nature 563, 203–208 (2018).

14. W. Shih, S. Yamada, N-cadherin as a key regulator of collective cell migration in a 3D environment. Cell Adh Migr 6, 513–517 (2012).

15. Y. Attieh et al., Cancer-associated fibroblasts lead tumor invasion through integrin-beta3-dependent fibronectin assembly. J Cell Biol, (2017).

16. S. Gopal et al., Fibronectin-guided migration of carcinoma collectives. Nat Commun 8, 14105 (2017).

17. B. Erdogan et al., Cancer-associated fibroblasts promote directional cancer cell migration by aligning fibronectin. J Cell Biol 216, 3799–3816 (2017).

18. R. Pankov, A. Momchilova, N. Stefanova, K. M. Yamada, Characterization of stitch adhesions: Fibronectin-containing cell-cell contacts formed by fibroblasts. Exp Cell Res 384, 111616 (2019).

19. P. Kollmannsberger, C. M. Bidan, J. W. C. Dunlop, P. Fratzl, V. Vogel, Tensile forces drive a reversible fibroblast-to-myofibroblast transition during tissue growth in engineered clefts. Sci Adv 4, eaao4881 (2018).

20. P. Cheung et al., Regenerative Reprogramming of the Intestinal Stem Cell State via Hippo Signaling Suppresses Metastatic Colorectal Cancer. Cell Stem Cell 27, 590–604 e599 (2020).

21. L. De Smedt, S. Palmans, X. Sagaert, Tumour budding in colorectal cancer: what do we know and what can we do? Virchows Arch 468, 397–408 (2016).

22. N. Ohike et al., Tumor budding as a strong prognostic indicator in invasive ampullary adenocarcinomas. Am J Surg Pathol 34, 1417–1424 (2010).

23. F. Prall, Tumour budding in colorectal carcinoma. Histopathology 50, 151–162 (2007).

24. H. C. van Wyk et al., The relationship between tumour budding, the tumour microenvironment and survival in patients with primary operable colorectal cancer. Br J Cancer 115, 156–163 (2016).

