## Extended Data for "Cancer-associated fibroblasts actively compress cancer cells and modulate mechanotransduction"

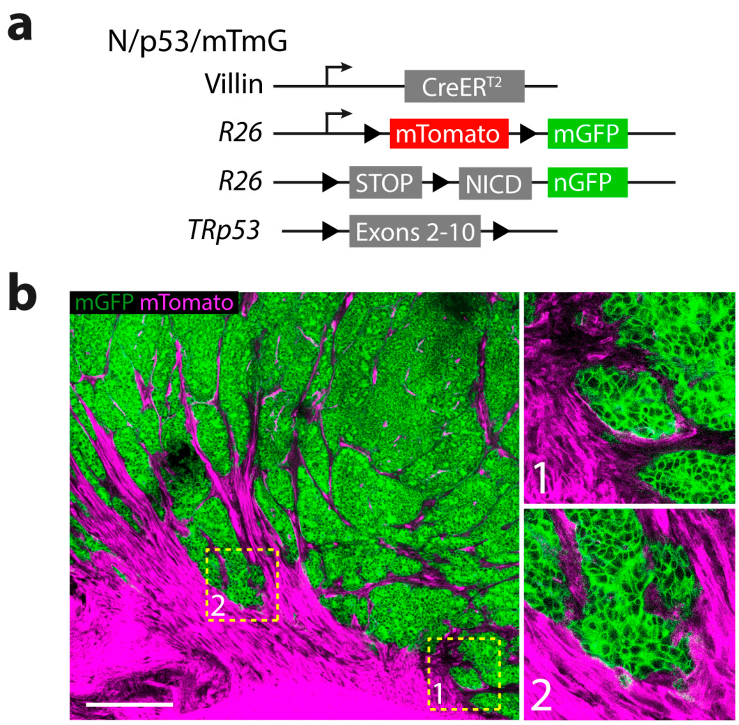


**Extended Data Fig. 1. CAFs form intratumoral capsules**

**a,** Schematic representation of the NICD/p53/mTmG mouse model. **b,** Representative image of a NICD/p53/mTmG intestinal tumor. Cancer cells (membrane-GFP, green), stromal cells (membrane-tdTomato, magenta). Scale bar, 300 µm. Right panels, magnified boxed regions 1 and 2.

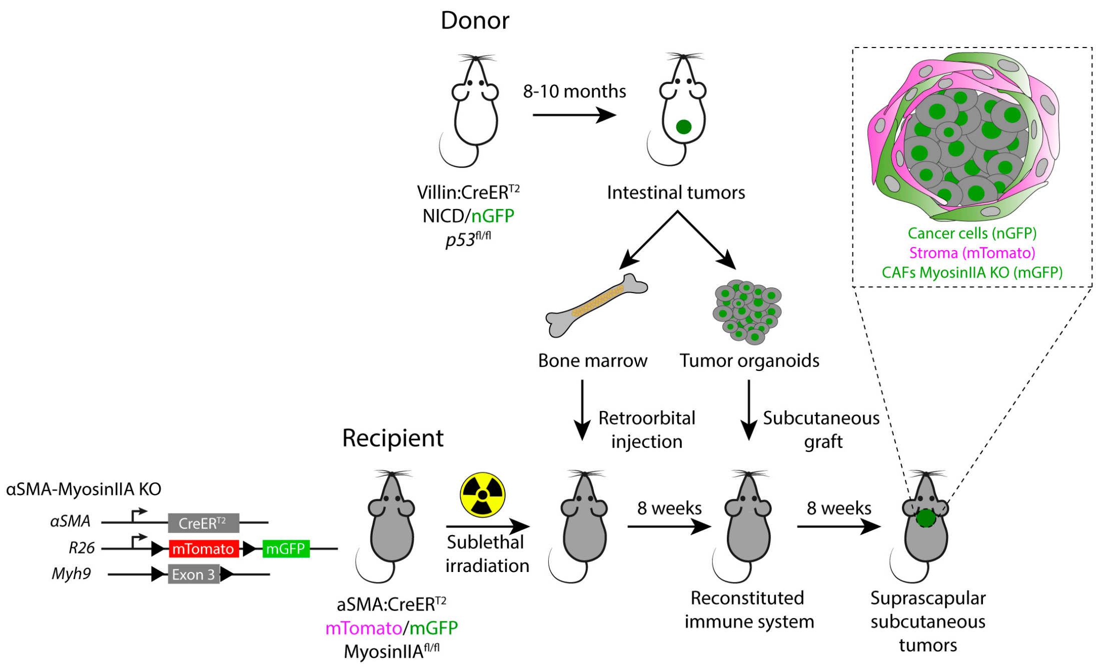


**Extended Data Fig. 2. Generation of subcutaneous tumors in αSMA-MyosinIIA KO mice**

Left, schematic representation of the αSMA-MyosinIIA KO mouse model. Right, experimental design for the engraftment of NICD/p53/mTmG tumor organoids in αSMA-MyosinIIA KO mice.

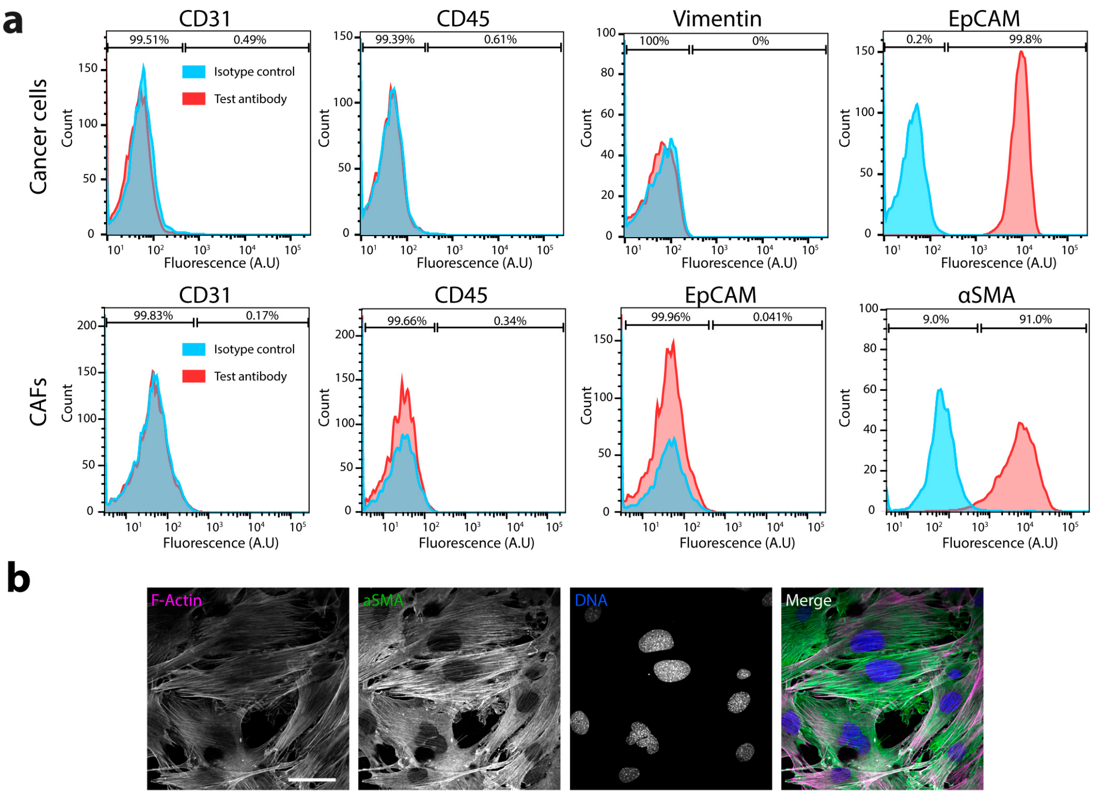


**Extended Data Fig. 3. Characterization of primary CAFs and cancer cells**

**a**, Flow cytometry analysis of the expression of CD31, CD45, Vimentin and EpCAM in primary cancer cells, and of CD31, CD45, EpCAM and αSMA in primary CAFs. The percentages (%) of stained cells (red) are plotted relative to the basal signal of an isotype control (light blue). **b**, Representative images of primary CAFs stained for F-Actin (phalloidin, magenta), αSMA (green) and DNA (DAPI, blue). Scale bar, 50 µm.

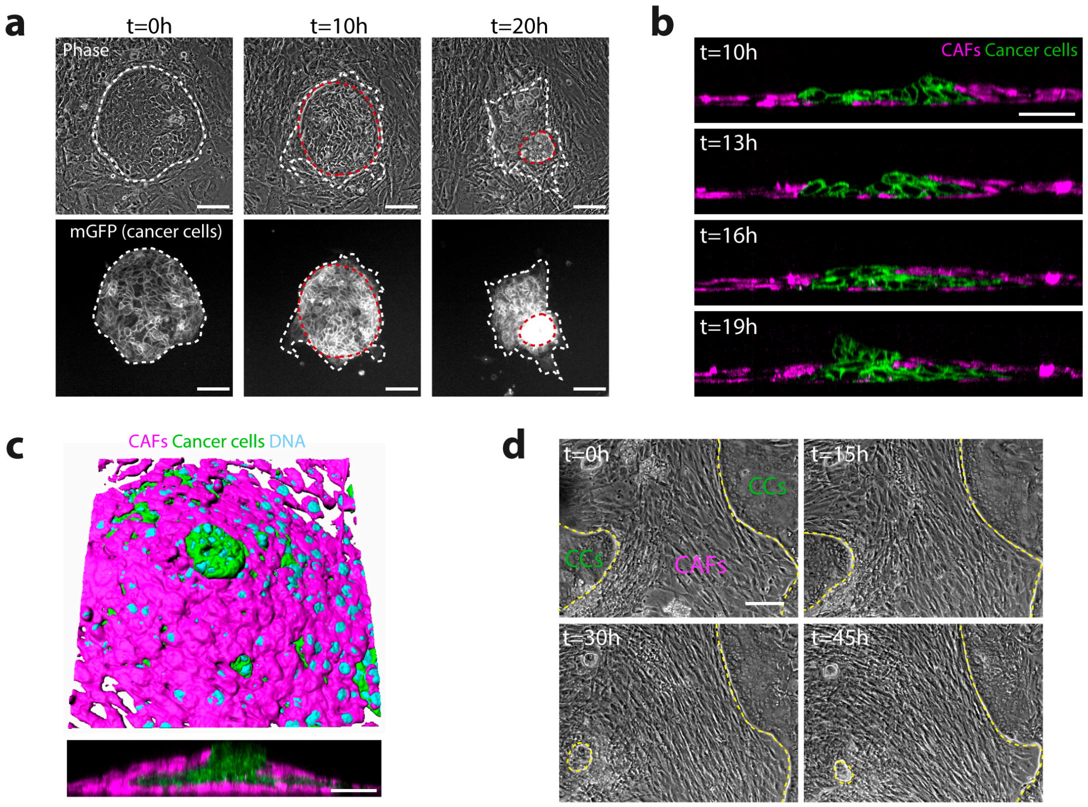


**Extended Data Fig. 4. CAFs encapsulate and deform N/p53 tumor cell clusters**

**a**, Representative images of CAFs cocultured with N/p53 tumor cells, evolving over time. Top panels, phase contrast. Lower panels, cancer cells (membrane-GFP, green) Red dashed lines delineate the contour of CAFs ring. White dashed lines delineate the contour of the tumor cell cluster. Scale bar, 100 µm. **b**, Representative orthogonal images of CAFs cocultured with N/p53 tumor cells, evolving over time. Cancer cells (membrane-GFP, green) and CAFs (CellTracker, magenta). Scale bar, 100 µm. **c**, 3D rendering of CAFs-N/p53 tumor cells co-cultures after 24h in culture. Cancer cells (membrane-GFP, green) and CAFs (CellTracker, magenta), DNA (DAPI, cyan). Lower panel, orthogonal Z section. Scale bar: 50µm. **d**, Representative images of CAFs and cancer cells that exited a PDX tissue fragment and spontaneously organized over time. Yellow dashed lines represent the CAFs edge and ring. Scale bar, 100 µm.

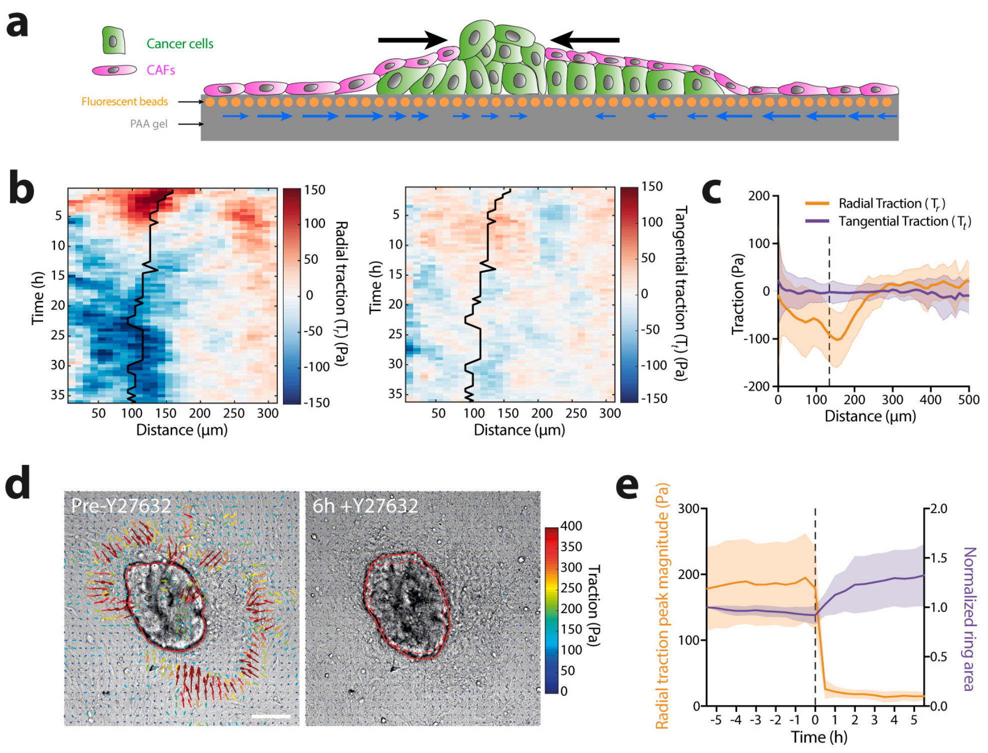


**Extended Data Fig. 5. CAFs compress cancer cells using actomyosin contractility**

**a**, Schematic representation of the interpretation of force patterns generated by CAFs during cancer cells deformation and budding. Contraction of CAFs rings generate inward pointing forces (black arrows) that are transmitted through the CAFs monolayer to the polyacrylamide (PAA) gel (grey), generating a pattern of inward pointing radial tractions (blue arrows). **b**, Representative kymograph of circumferentially averaged radial (left) and tangential (right) tractions as a function of the distance to the center of the cancer cell cluster. Solid black line represents the cancer cell cluster contour. **c**, Circumferentially averaged tangential tractions $T_t$ (purple) and radial tractions $T_r$ (orange) as a function of the distance to the cancer cell cluster center after 36h of co-culture. Black dashed line represents the cluster boundary. Data represented as mean ± SD, n=49 clusters from N=3 independent experiments. **d**, Traction maps overlaid on a DIC image of cancer cell-CAFs after ≈40h co-culture (Pre-Y27632) and 6h after addition of Y27632 (6h+Y27632). Solid red line represents the contour of CAFs ring; dashed line represents the contour of CAFs ring before Y27632 addition. Scale bar, 100 µm. **e**, Circumferentially averaged radial traction peak magnitude near the boundary of the cancer cell cluster (orange) and CAFs ring area (purple) normalized to the initial ring size, as a function of time. Dashed black line represents the moment of Y27632 addition. Data represented as mean ± SD, n=30 clusters from N=3 independent experiments.

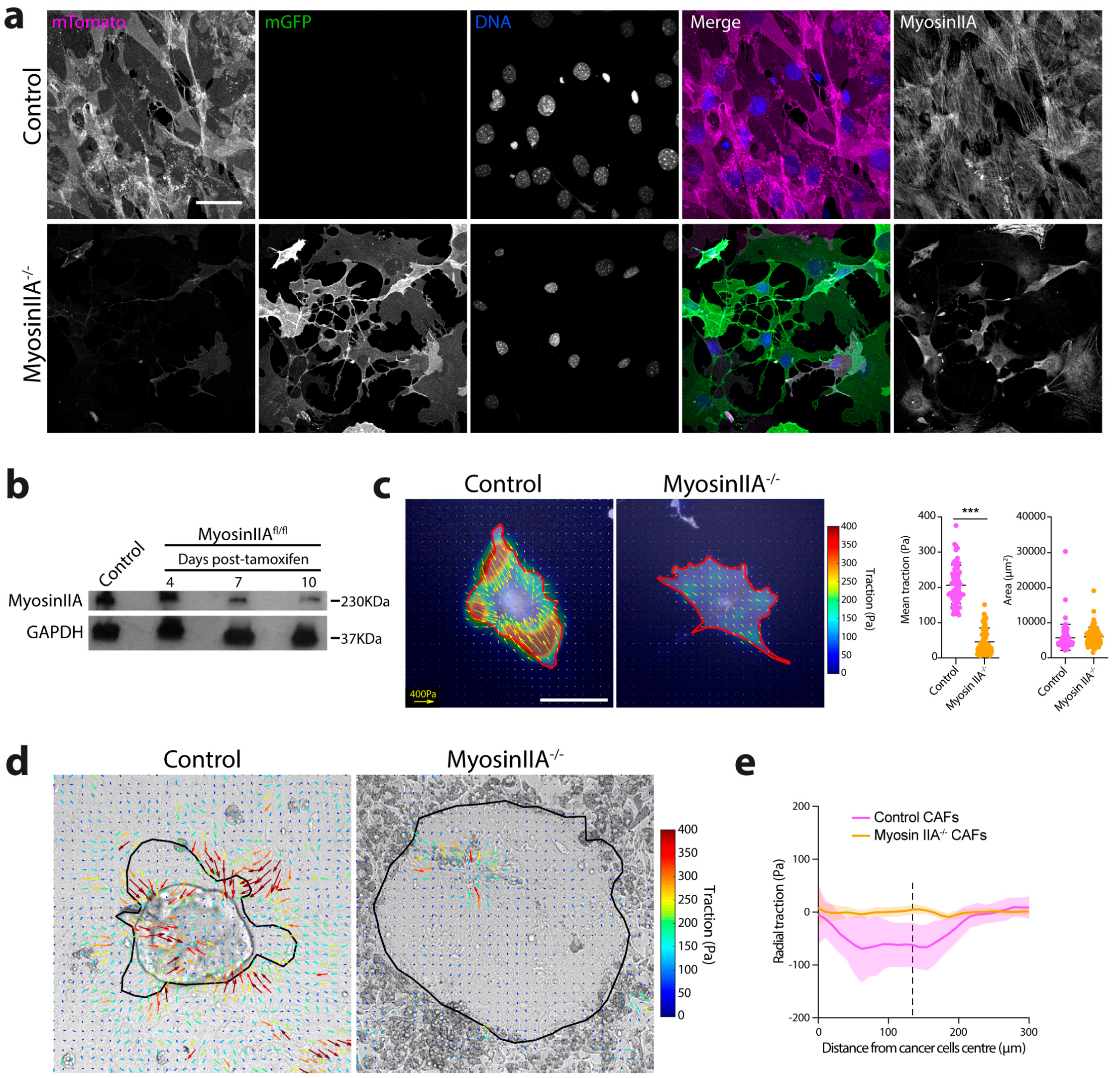


**Extended Data Fig. 6. MyosinIIA KO CAFs do not exert traction forces or deform cancer cells**

**a**, Representative images of control (upper panels) and MyosinIIA KO (lower panels) CAFs after 7 days of in vitro knockout induction with 4-hydroxytamoxifen. Control CAFs (membrane tdTomato, magenta), KO CAFs (membrane-GFP, green, and low levels of membrane-tdTomato, magenta). Cells were stained for DNA (DAPI, blue) and MyosinIIA (right panels). Scale bar, 50 µm. **b**, Western blot for MyosinIIA (230KDa) in control and MyosinIIA KO CAFs after 4, 7 and 10 days of knockout induction. GAPDH (37KDa) was used as loading control. **c**, Representative traction maps overlaid on images of control (membrane-tdTomato) and MyosinIIA KO (membrane-GFP) single CAFs after 7 days of in vitro knockout induction. Solid red line represents the cell contour. Scale bar, 50µm. Scale vector, 400 Pa. Right dot plots, quantification of mean traction and area per cell. Data represented as mean ± SD. n=75 cells, from N=3 independent experiments. *** $p<0.001$, Mann Whitney test. **d**, Representative traction maps overlaid on a DIC image of a cancer cell and control (left) or myosin IIA KO (right) CAFs after 48h of culture. Black line represents the contour of the cancer cell cluster Scale bar, 100 µm. **e**, Circumferentially averaged radial tractions $T_r$ in co-cultures of cancer cells and control (magenta) and MyosinIIA KO (orange) CAFs after 48h, as a function of the distance to the cancer cell cluster center. Black dashed line represents the cluster boundary. Data represented as mean ± SD. Control, n=36 clusters, MyosinIIA KO, n=30 clusters, from N=2 independent experiments.

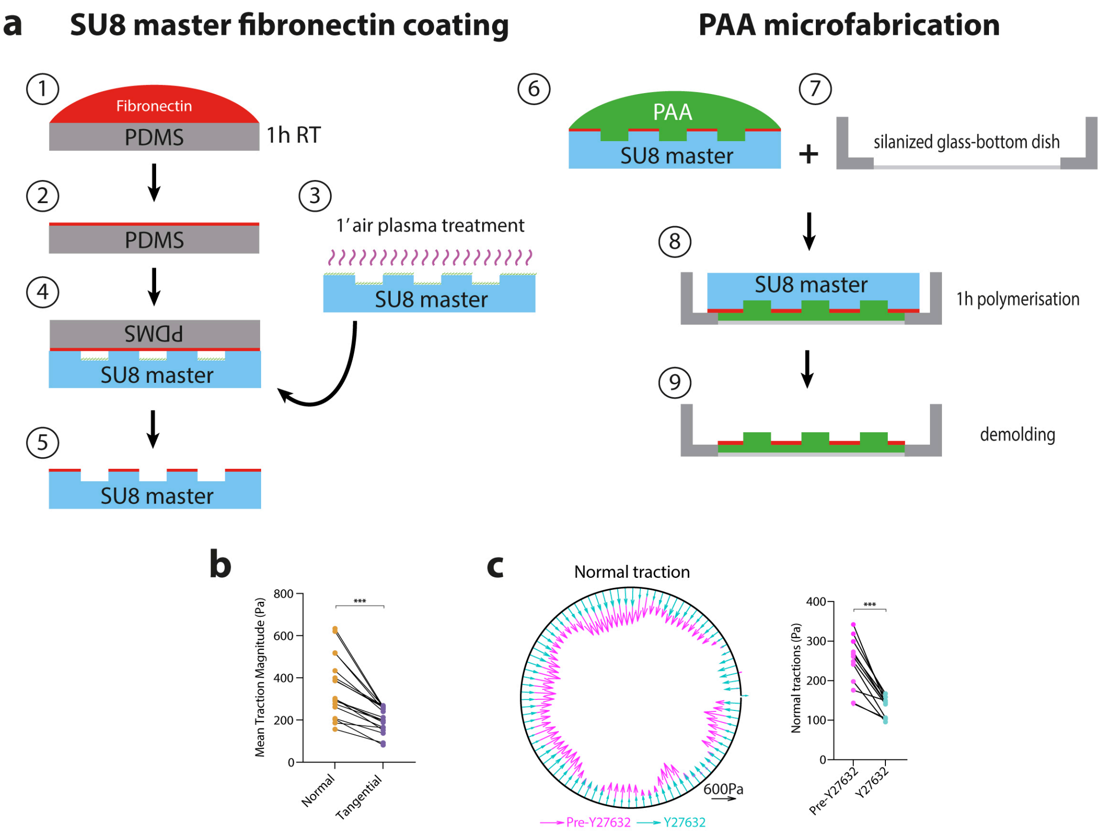


**Extended Data Fig. 7. Pillar compression depends on CAFs contractility**

**a**, Schematic representation of the methodological approach for pillar microfabrication. **b**, Mean normal and tangential traction magnitude in control pillars. n=18 pillars, from N= 3 independent experiments. *** p<0.001, Wilcoxon matched-pairs signed rank test. **c**, Representation of CAF normal tractions averaged across pillar height, on a representative pillar before (magenta) and after (blue) Y27632 treatment. Scale vector, 600 Pa. Right dot plot, quantification of mean normal traction for each pillar before and after Y27632. n=12 pillars, from N=2 independent experiments. *** p<0.001, Wilcoxon matched-pairs signed rank test.

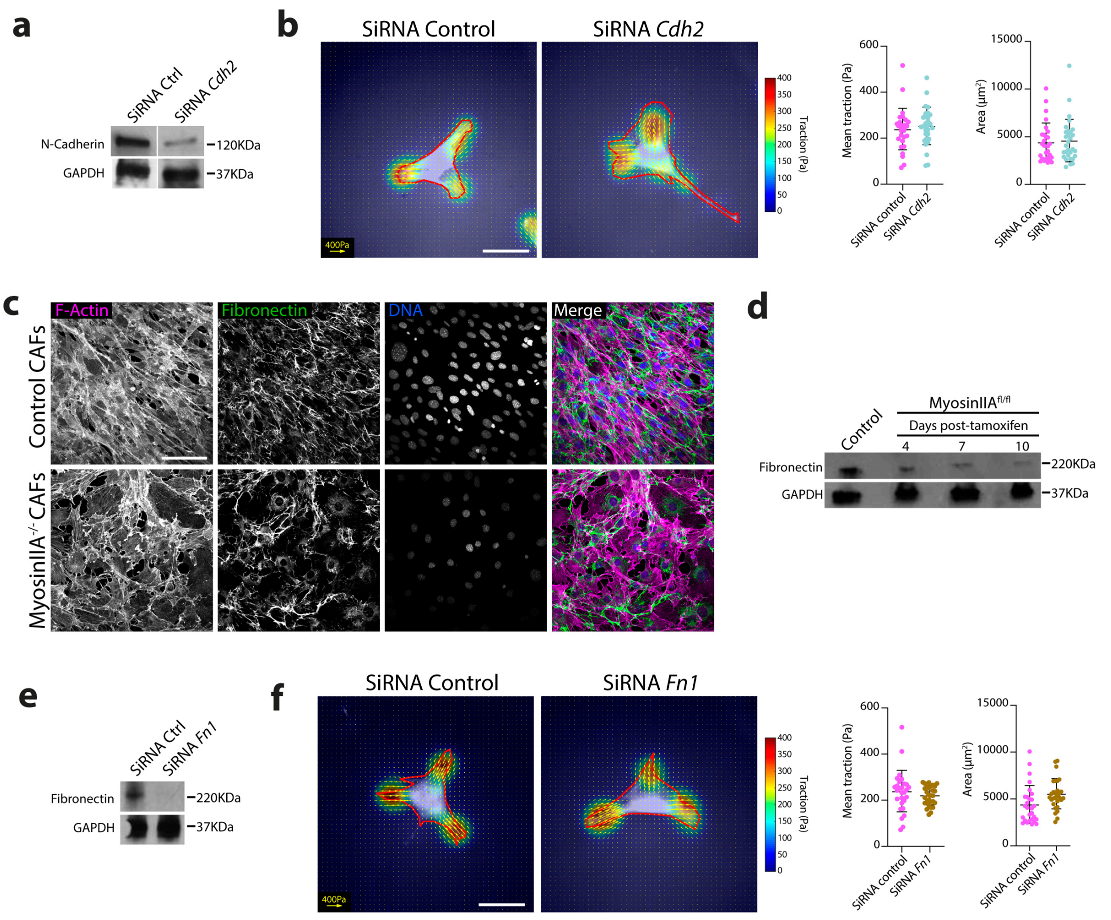


**Extended Data Fig. 8. Fibronectin and N-cadherin knockdown does not affect force generation at single cell level**

**a**, Western blot for N-Cadherin in control and SiRNA Cdh2 CAFs. GAPDH was used as loading control. **b**, Representative traction maps overlaid on phase contrast images of control and SiRNA Cdh2 single CAFs. Solid red line represents the cell contour. Scale bar, 50 µm. Scale vector, 400 Pa. Right dot plots, quantification of mean traction and area per cell. Data represented as mean ± SD. Control, n=28 cells, SiRNA Cdh2, n=30 cells, from N=2 independent experiments. Non-significant, Mann Whitney test. **c**, Representative images of control (upper panels) and MyosinIIA KO (lower panels) CAFs, after 7 days of in vitro knockout induction with 4-hydroxytamoxifen. Cells were stained for F-actin (Phalloidin, magenta), Fibronectin (Green) and DNA (DAPI, blue). Scale bar, 100µm. **d**, Western blot for Fibronectin in control and MyosinIIA KO CAFs after 4, 7 and 10 days of knockout induction. GAPDH was used as loading control. **e**, Western blot for Fibronectin in control and SiRNA Fibronectin CAFs. GAPDH was used as loading control. **f**, Representative traction maps overlaid on phase contrast images of control and fibronectin-KD CAFs. Solid red line represents the cell contour.

Scale bar, 50 µm. Scale vector, 400 Pa. Right dot plots, quantification of mean traction and area per cell. Data represented as mean ± SD. Control, n=28 cells, SiRNA Fn1, n=29 cells, from N=2 independent experiments. Non-significant, Mann Whitney test.
