## Supplementary Data for "Cancer-associated fibroblasts actively compress cancer cells and modulate mechanotransduction"

**This PDF file includes:**

Materials and Methods

Captions for Supplementary Videos 1 to 11

Supplementary Note 1

**Other Supplementary Materials for this manuscript include the following:**

Supplementary Videos S1 to S11

#### Materials and Methods

##### Mouse models

For animal care, we followed the European and French National Regulation for the Protection of Vertebrate Animals used for Experimentation and other Scientific Purposes (Directive 2010/63; French Decree 2013-118). The project was authorized by the Animal Welfare Body at Institut Curie Research Centre APAFIS#25603-2020053122444776. All mice were kept under Specific Pathogen Free (SPF) conditions for breeding.

Double fluorescent pVillin-CreER$^{T2}$, LSL-NICD-nGFP; mT/mG; p53$^{fl/fl}$ mice were generated as previously described[10]. Four-weeks old mice were injected intraperitoneally with tamoxifen (50 mg/kg) for 5 consecutive days for induction of Cre recombinase activity. Approximately 8 months after tamoxifen injection mice spontaneously develop invasive intestinal carcinomas[25]. All cells from these mice express a membrane-targeted tdTomato, while tumor cells express a nuclear-targeted GFP.

Double fluorescent SMA-CreER$^{T2}$; R26$^{mT/mG}$; myosin IIA$^{fl/fl}$ were generated by crossing mice containing the MyoIIA heavy-chain (*Myh9*) floxed[26] with mice expressing Cre recombinase under the control of the SMA promoter[27], and with Rosa26-mTmG[28] mice. Rosa26-mTmG mice express a membrane-targeted tandem dimer Tomato (mT) prior to Cre-mediated excision and a membrane-targeted GFP (mG) following excision. Membrane targeting was achieved using the MARCKS membrane tag[28].

##### Generation and culture of tumor organoids

Tumors from pVillin-CreER$^{T2}$, LSL-NICD-nGFP; mT/mG; p53$^{fl/fl}$ mice were excised and dissociated using a scalpel in medium containing DMEM-F12 (ThermoFisher Scientific) supplemented with 2,5% (v/v) GlutaMAX (Gibco), 2% (v/v) Antibiotic-Antimycotic (Gibco) and 300 units/ml of Collagenase III (StemCell). Tissue pieces were incubated for 2 h at 37°C under agitation (180 rpm), and then filtered first through 100 µm and 40 µm filters. Dissociated cells were centrifuged and resuspended in a 100 µl drop containing a mix of 50% Matrigel (Corning)-50% (v/v) tumoroid medium (DMEM-F12, supplemented with 2,5% (v/v) GlutaMAX, 2% (v/v) antibiotic-antimycotic, 100ng/mL Noggin (Peprotech), 50 ng/mL EGF (Peprotech), 10 ng/mL (Peprotech), 1% (v/v) B27 supplement (ThermoFisher Scientific) and 1% (v/v) N-2 supplement

(ThermoFisher Scientific)). Matrigel drops were allowed to polymerize for 40 mins at 37°C and 5% $CO_2$ and were then covered with 2 mL of tumoroid medium. Once formed, tumoroids were split once a week.

**Tumor establishment in αSMA MyoIIA KO mice**

Bone marrow transplantation experiments were performed in order to render the immune system of αSMA MyoIIA KO mice compatible with tumor growth.

*Irradiation.* 6-8 weeks old recipient mice (SMA-CreER$^{T2}$; R26$^{mT/mG}$; myosin IIA$^{fl/fl}$) were placed in an acrylic container in continuous airflow between two opposite X-ray sources (CIXD Dual Irradiator, Xstrahl). Cre$^{-/-}$ mice were used as controls. Mice were exposed to a Fractioned Total Body Irradiation (FTBI), at a rate of 1,18 Gy/min for a total dose of 10 Gy, fractionated in two 5 Gy doses with a 4 h interval.

*Bone marrow transplantation.* Donor mice (non-induced pVillin-CreER$^{T2}$, LSL-NICD-nGFP; mT/mG; p53$^{fl/fl}$, 4-10 months old) were sacrificed and bone marrow cells (BMCs) were collected from the tibias, femurs and humerus, and resuspended in 300 µl of PBS containing 2% (v/v) FBS. Immediately after irradiation, sex-matched recipient mice were injected retro-orbitally with 100 µl of the BMCs suspension, containing approximately 2x10$^6$ BMCs, using a 27-gauge needle. Mice were left for 8 weeks to allow efficient grafting of BMCs and reconstitution of the immune system.

*Tumor establishment and MyosinIIA KO induction.* Tumoroids from pVillin-CreER$^{T2}$, LSL-NICD-nGFP; mT/mG; p53$^{fl/fl}$ mice were cultured as described above. Prior to injection in mice, tumoroids from 12 24-well Matrigel drops were harvested and mechanically disaggregated using a pipette tip, centrifuged and resuspended in 100 µl of 1:1 Matrigel/culture medium. This solution was injected into the interscapular fat pad of bone marrow transplanted mice. For this, under Isoflurane gas anesthesia, a small skin incision was performed at the level of the interscapular region, the fat pad was exteriorized and tumoroids were injected directly into it using a 25 G needle. Mice were then sutured using wound clips (7.5*1.5 mm, Phymep), and placed into standard housing conditions during the time of tumor development. 3 weeks post engraftment mice were injected daily with tamoxifen (50 mg/kg) for two consecutive days, to induce Cre-mediated gene recombination. Tamoxifen was re-administered either every week or every other week throughout the entire duration of the experiment to ensure efficient knockout of possibly newly generated CAFs. For tumor compartmentalization analyses we used mice injected every week with tamoxifen. For laser ablation experiments, all mice were analyzed, as ablations were performed specifically in areas with high content of knockout CAFs ($mGFP^{+}$) To account for possible secondary effect of tamoxifen in tumor development, all mice ($Cre^{+/-}$ and $Cre^{-/-}$ controls) were injected with the same doses of tamoxifen. 9 weeks after tumoroid injection mice were sacrificed, tumors were excised and prepared for downstream analysis.

**Immunostaining if tumor tissue sections**

Tissue was fixed in 4% paraformaldehyde (Electron Microscopy Sciences)/PBS (v/v) for 20 mins at RT, and then dehydrated first in 15% sucrose (Sigma-Aldrich)/PBS (w/v) solution for 1h and then in 30% sucrose/PBS (w/v) solution for 2h at RT. After, tissue was embedded with OCT compound (Sakura) in plastic gaskets (Euromedex), frozen at -20°C and cut on the cryostat using SuperFrost Plus™ Adhesion slides (VWR, Menzel Gläser). Tissue sections were then permeabilized with 0.2% Triton x100 (Sigma-Aldrich)/PBS (v/v) for 1h at RT, blocked with 3% BSA (w/v) (IgG-Free, Protease-Free, Jackson Immuno Research) in 0.05% Triton x100/PBS (v/v) solution for 1h at RT and stained with primary antibodies overnight in humidified chambers at RT. Sections were then washed 3 times with 0.05% Tx100/PBS (v/v) solution for 1h, incubated with secondary antibodies, DAPI and phalloidin, depending on the staining, for 4h at RT, washed 3 times in 0.05% Tx100/PBS (v/v) solution for 1h and mounted using AquaPolyMount (Polysciences). Antibodies references and dilutions are listed in Supplementary Table 1.

**Whole-mount staining of tumor tissue**

Fixed tissue was sliced to 250 µm thick slices in a vibratome (Leica VT1200S) as described before[10,29]. Staining was performed as described above with minor modifications: permeabilization was done with 1% Triton X-100/PBS (v/v) for 1h at RT (500 µl per tube), all antibody solutions and washing steps were performed using 0.2% Triton X-100/PBS (v/v), under mild shaking conditions (1 mL per tube) and concentration of antibodies was increased (see supplementary table 1, in 150 µl per tube). Incubations with antibodies were done without agitation.

**Confocal imaging**

Stained cryosections and whole-mount tissue, as well as stained in vitro cell cultures were imaged using an inverted confocal microscope (Zeiss LSM880NLO) using laser lines 405, 488, 561 and 633 nm and the following objectives: 25×/0,8 OIL, W, Gly LD LCI PL APO (UV) VIS-IR, 40x/1.30 OIL DICII PL APO (UV) VIS-IR and 63x/1.4 OIL DICII PL APO. Images were processed using ImageJ.

Second harmonic imaging of non-stained whole-mount tissue sections was performed in an inverted AOBS two-photon laser-scanning confocal microscope (Leica SP8), coupled with a femtosecond laser (Chameleon Vision II, Coherent Inc.) using a 40x/1.10 HC PL APO CS2 water immersion objective. The excitation wavelength was set at 947 nm and signals were acquired using 3 non-descanned HyD detectors: 525/40 nm (for GFP), 585/40 nm (for tdTomato) and <492 nm (for SHG). Acquisition was performed in resonant mode, with a Z-step of 5 µm and tiling arrays were performed in order to image large tissue areas. Image stitching was performed using the LAS X software (Leica).

**Segmentation of SHG images to quantify tumor compartmentalization.**

Tumor slices are generally tilted and folded, hindering the selection of one single plane of the Z-stack to perform 2D image analysis. Conventional maximum intensity or mean intensity projections fail at preserving low intensity regions, which are important to define the areas devoid of stroma. To overcome this, we performed Z-projections based on the intensity of big areas, not single pixels, in order to preserve the low intensity pixels. To achieve this, we calculated the local most in-focus plane (defined as the plane with maximum standard deviation) along the image using a scanning window of 300x300 pixels with an overlap of 0.33. This protocol provided a matrix with the number of the best focus plane in each of these 300x300 regions. We resized this matrix to the size of the original image and smoothened it by applying a Gaussian blur (600 pixels radius). This final matrix of Z-planes was used to perform projections of the Z-stacks and build 2D images.

To quantify areas devoid of stroma, we first applied a Gaussian blur (3 pixels radius) to the projected SHG images and applied a user-defined threshold to obtain a binary image labelling all the pixels lacking SHG signal (stromal devoid). We dilated and eroded (3 pixels) this image to fuse adjacent objects and indexed each independent object in the field, which was then filled and smoothened. A stromal devoid region was defined as any object bigger than 15565 $\mu m^2$ (circle of ~70 µm radius).

**Laser ablations**

CAFs-cancer cells cocultures were imaged using a two-photon laser-scanning microscope (Zeiss LSM880NLO) in single photon mode, using a 40x/1.30 OIL DICII PL APO (UV) VIS-IR objective, and laser lanes 488 and 561. CAF rings were ablated using a Ti:Sapphire laser (Mai Tai DeepSee, Spectra Physics) set at 800nm and laser power of 10%  Image acquisition was started 10 s before the ablation, every 2 s and for a total time of 50 s.

To perform laser ablations *in vivo,* living tumor tissue slices from control or myosinIIA KO mice were prepared as described previously[10,29], and placed into a 35 mm glass-bottom culture dish. A slice anchor (SHD-26GH/10; Harvard Apparatus) was placed on top of the tissue slices to minimize sample drift and covered with a drop (100µl) of DMEM-GlutaMAX (Glibco), supplemented with 1% (v/v) antibiotic-antimycotic (Gibco), 2,5% (v/v) fetal bovine serum, 1% (v/v) Insulin-Transferrin-Selenium (ITS, ThermoFisher Scientific) and 10 ng/mL EGF (Peprotech). The ablation setup was adapted from the in vitro experiments, with laser power set at 20%. Image acquisition was started 10 s before the ablation, every 2 s and for a total time of 50 s. Para-Nitroblebbistatin (50 µM) (Optopharma) was added to 4-5 slices per mouse, incubated at 37°C, 5% $CO_2$ for 2.5 h, and then ablations were performed as described.

**Quantification of tissue recoil**

To quantify tissue recoil upon laser ablation, we measured tissue displacements using custom-made PIV with a window size of 16x16 pixels and an overlap of 0.75. For each timepoint, we computed tissue displacements relative to the timepoint immediately before ablation.

We measured displacements of tdTomato and GFP channels independently. Displacements of each channel were combined according to the segmentation of different cell populations. For all conditions, cancer cell regions (GFP) were manually segmented. For in vitro and in vivo control tumors, this segmentation was enough, since CAFs/stroma express tdTomato. For myosin-IIA KO tumors where the stroma has a heterogeneous labelling, threshold-based automatic segmentation was performed to discriminate between KO CAFs (GFP+, mTomato) and the rest of the stroma (GFP-, mTomato+). Based on this segmentation, we build a combined displacement map containing, at each pixel, the displacements of the relevant channel.

To quantify the recoil of cancer cells and CAFs near the cut, we define different regions of interest (ROIs) where displacements will be averaged. We automatically define these regions by dilating the mask of the ablated region 48 pixels for cancer cells and 75 pixels for CAFs, and then finding the overlap between these dilated regions and the cancer cell or the stromal area, respectively. Control ROIs far from the cut were defined by automatically creating ROIs within the cancer cells as far as possible from the cut. We then averaged displacements in each of these ROIs and computed the magnitude and direction of the mean displacement. For ablations parallel to the cancer cell boundary where the cut is not a straight line, we decomposed displacements in perpendicular and tangential components respect to the cut (as explained above for traction forces) before averaging. We used the mean perpendicular and tangential components to calculate the angle respect to the cut defined over 360º (-90º towards the cut, 0 or 180º as parallel to the cut, and 90º away from the cut). For laser ablations perpendicular to the cancer cell boundary, the direction of the cut was defined by fitting a straight line to the cut. The angle difference between the mean displacements and the cut was then calculated (-90º towards the cut, 0 or 180º as parallel to the cut, and 90º away from the cut for CAFs and 0º towards the cut, +90 or -90º as parallel to the cut, and 180º away from the cut for the cancer cells).

**Quantification of YAP localization**

YAP localization was assessed by quantifying the 3D correlation between Z-stacks of DAPI and YAP channels of immunostainings. First, cell nuclei were segmented in each plane of the Z-stack using a user-defined threshold. For each plane, the masks of the nuclei were dilated 5 pixels to also include cytoplasmic areas. The rest of the pixels were set to not a number, thus excluding cell-free areas from the analysis. Pearson correlation coefficient between YAP and DAPI images was measured using a 3D scanning window of 32x32x3 pixels (xyz) with an overlap of 0.75. The window evaluates the correlation along the XY area, thus providing 2D matrices of YAP nuclear correlation. For in vitro images, the window was centered around the plane of maximum

intensity in the DAPI channel. For in vivo images, where the tissue is not flat, the scanning window was also allowed to move in Z to center around the plane of maximum local intensity.
For in vitro images, cancer cells were manually segmented to calculate their average YAP nuclear correlation. For in vivo images, which are heterogeneous and contain multiple cell types, cancer cells were automatically segmented using a threshold based on their fluorescence profile: no mTomato and low but positive GFP (to exclude GFP+ CAFs in MyosinIIA KO tumors).

**Patient derived xenografts (PDX)**

Tumor tissue was obtained from rectal cancer patients at the moment of surgery after chemo-radiotherapy treatment at Institut Curie Hospital, Paris, with the patients' written consent and approval of the local ethics committee. Samples were collected in DMEM with 10 mmol/L HEPES, 4.5 g/L glucose, 1 mmol/L pyruvate sodium, 200 U/mL penicillin, 200 µg/mL streptomycin, 5 µg/mL ciprofloxacin, 20 µg/mL metronidazole and 2.5 µg/mL fungizone. Small fragments (~50 $mm^3$) were subcutaneously engrafted into the scapular area of anesthetized Nude mice (Crl:NU(Ico)-$Foxn1^{nu}$) (either under xylazine/ketamine or isoflurane anesthesia). Tumor growth was monitored weekly, and tumors were excised when reached a volume between 800 to 1500 $mm^3$ and passaged into a new recipient mouse. PDXs were considered as established after 3 consecutive passages.

**Primary CAF lines generation and characterization**

Primary mouse CAFs were obtained either from PDX tumor tissue or from tumors generated in SMA-$CreER^{T2}$; $R26^{mT/mG}$; myosin $IIA^{fl/fl}$ mice. For this, mice were sacrificed and tumors excised and collected in DMEM Glutamax with 2% (v/v) Antibiotic-Antimycotic (Gibco), 5 µg/mL ciprofloxacin and 20 µg/mL metronidazole, on ice. The tissue was sliced into small pieces of less than 1$mm^2$ and carefully plated on top of an 11 kPa polyacrylamide gel coated with 100 µg/ml of collagen I (see gel preparation protocol). Tissue pieces were then cultured in DMEM GlutaMAX supplemented with 10% (v/v) FBS, 1% (v/v) Insulin-Transferrin-Selenium (ThermoFisher Scientific), 2% (v/v) Antibiotic-Antimycotic (Gibco), and Metronidazol/Ciprofloxacin (20 and 5 µg/mL, respectively). Medium was changed every 3 days. Once emerged from the tissue, and while still being in the PAA gels, CAFs were immortalized by retroviral infection of SV40 large T-antigen as described[30]. pBABE-puro SV40 LT was a gift from Thomas Roberts (Addgene plasmid # 13970; http://n2t.net/addgene:13970; EEID: Addgene_13970)[31]. After immortalization cells were cultured in DMEM GlutaMAX supplemented with 10% FBS and 1% Insulin-Transferrin-Selenium (ThermoFisher Scientific).

Immortalized CAF cultures were characterized for the expression of αSMA, by immunofluorescence and flow cytometry, where CD45, EpCAM and CD31 were used as markers to exclude the presence of immune, epithelial or endothelial cells, respectively. See table 1 for antibody information and dilutions.

In vitro recombination of CAFs coming from SMA-CreER$^{T2}$; R26$^{mT/mG}$; myosin IIA$^{fl/fl}$ mice was induced using 2μM 4-hydroxytamoxifen (Sigma Aldrich), for 2 days.

**Establishment of human primary cancer cell lines and characterization**

Primary human cancer cells were obtained from PDX tumors. For this, mice were sacrificed, and tumors were excised and collected in DMEM GlutaMAX with antibiotics (same as above) on ice. Tumors were cut into 0,5 $cm^3$ blocks and crushed with a striated plunger from a disposable syringe. The resulting pieces were transferred to a 25 $cm^2$ culture flask and incubated in DMEM GlutaMAX, 10% (v/v) FBS, with the same antibiotics at 37°C, 5% $CO_2$. Antibiotics were removed after passage 2. If necessary, controlled trypsinizations were done to remove contaminating fibroblasts. Cancer cells were routinely passed once a week, and the medium was changed twice in between.

Primary cancer cell cultures were characterized by the expression of EpCAM by flow cytometry. CD45, Vimentin and CD31 were used to exclude the presence of immune, fibroblastic or endothelial cells, respectively.

**siRNA**

For protein (fibronectin and N-cadherin) depletions using siRNA, CAFs were cultured in standard conditions and transfected using Lipofectamine 3000 (ThermoFisher Scientific). $50x10^4$ CAFs were plated overnight in 6-well plates and then subjected to transfection using 125nM siRNA. Cells were used 48 h after transfection. For cancer cells-CAFs co-cultures experiments, a second transfection round was performed after co-culture establishment, right before image acquisition, to maximize knockdown efficiency. The specific siRNAs used were: Mouse Cdh2:

SI00168252 (Qiagen), Mouse Fn1: J-043446-09-0002 (Horizon Discovery, Perkin Elmer), Negative control: AllStars Negative Control siRNA (1027280, Qiagen).

**Western blot**

Cells were scrapped and lysed in RIPA buffer (10 nM Tris HCl pH 7.4, 100 mM NaCl, 1 mM EDTA, 1 mM EGTA, 1% (v/v) Triton X100, 10% (v/v) glycerol, 0,1% (v/v) SDS) with a 1% (v/v) cocktail of protease inhibitors (Sigma-Aldrich). Protein lysates were resolved on 4-20% or 7,5% TGX gels (Miniprotean TGX, BioRad), transferred onto nitrocellulose membranes (Blot Turbo, BioRad) and immunoblotted with the indicated antibodies (Supplementary Table 1) overnight at 4°C in 5% (w/v) non-fat milk in PBS-Tween (0,1% v/v), and detected using peroxidase-conjugated secondary antibodies (Supplementary Table 1), incubated for 1h at RT in 5% (w/v) non-fat milk in PBS-Tween (0,1% v/v). Signal was revealed using an ECL substrate (Thermo Pierce ECL 2) and visualized using X-ray films.

**Fabrication of polyacrylamide gel substrates**

Glass-bottom dishes (World Precision Instruments) were treated with silane (3-(Trimethoxysilyl) propyl methacrylate, Sigma-Aldrich) diluted 1:3 in PBS, for 15 mins, rinsed 3 times (5 mins each time) with water, and dried out. They were then incubated with a solution of 0,5% Glutaraldehyde/PBS (w/v) for 30 mins, rinsed 3 times with water and dried out. Polyacrylamide (PAA) gels of 11kPa (Young modulus) were produced as described previously[32]. Briefly, a solution containing 7,5% (v/v) acrylamide, 0,1% (v/v) bis-acrylamide, 0,5% (w/v) ammonium persulphate, 0,05% (w/v) tetramethylethylenediamine and 2% (v/v) of 200-nm-diameter red/green/DAPI fluorescent carboxylate-modified beads was prepared and allowed to polymerize on top of the silanized glass, covered with a non silanized 18-mm-diameter coverslip. Beads were not added for gels used to generate primary CAF cultures. After 1 h, the coverslip was removed and the PAA gel surface was incubated with a solution of 2 mg/mL Sulpho-SANPAH under a 265 nm UV light for 10 minutes. After that, gels were washed once with 10 mM HEPES for 3 minutes under agitation, and twice with PBS, to remove the excess of Sulpho-SANPAH. Gels were then coated with a solution of 100 µg/mL rat-tail Collagen I (Corning) in PBS overnight at 4°C.

**Fabrication of PDMS stencils**

Polydimethylsiloxane (PDMS) membranes were fabricated as explained previously[32]. Briefly, SU8-50 masters containing an array of 150 µm radius circles were raised using conventional photolithography. Uncured PDMS was spin-coated on top of the masters to a thickness lower than the SU8 features (35 µm) and cured at 80 °C for 2 hours. A thick border of PDMS was added for handling purposes. Finally, PDMS stencils were peeled off and stored in 96% ethanol until use.

**Cell patterning on PAA gels**

PDMS stencils were treated with a solution of 2% Pluronic acid F127/PBS (w/v) (Sigma Aldrich) for one hour. They were then rinsed twice in PBS, let dry for 20 mins and carefully placed on top of a well-dried, collagen-coated PAA gel. ~500.000 cancer cells were seeded in a 100 µl drop on top of the PDMS stencil. After 2 h, non-attached cells were removed, and 2 mL of medium supplemented with 10% (v/v) FBS, 2% (v/v) Antibiotic-Antimycotic (Gibco), and Metronidazole/Ciprofloxacin (20 and 5 µg/mL, respectively) was added. The attached cancer cells were then allowed to spread overnight. PDMS stencils were then carefully removed and a cell culture medium containing the same antibiotics was added.

If necessary, cancer cells were labeled at this step. For this, the culture medium was removed, and cells were washed twice with PBS and incubated with a solution of CellTracker Orange (ThermoFisher Scientific) (1:1000 in Hanks' Balanced Salt Solution, HBSS, Gibco), for 30 mins at 37°C, 5% $CO_2$. Cells were then washed three times (once with HBSS and twice with cell culture medium), for 10 minutes each. If necessary, CAFs were as well stained, before being detached from the culture plate. For this, CAF cultures were incubated with a solution of CellTracker Green (ThermoFisher Scientific) (1:2000 in culture medium without FBS) for 30 mins at 37°C, 5% $CO_2$. After staining, CAFs were washed 3 times (10 minutes each) with a complete culture medium.

To generate cancer cells-CAFs cocultures, the culture medium was removed from the plate containing patterned cancer cell cultures, and a solution containing ~$2.5x10^5$ CAFs in a 75 µl drop was added on top of the PAA gel surface. Cells were then incubated for 30-60 minutes to allow CAFs to attach to the gel. Non-attached cells were rinsed out with cell culture medium, and 3 mL of medium supplemented with antibiotics (same as above) were added. Cocultures were incubated at 37°C, 5% $CO_2$, or immediately used for experiments.

**Traction force microscopy**

Time-lapse images were acquired on an automatic inverted microscope (Nikon Eclipse Ti-E) equipped with thermal, $CO_2$, and humidity control, using a 10X objective (NA 0.5 DIC, air) and controlled through MetaMorph (Universal Imaging). 10-20 cancer cell islands per experimental condition were imaged every 30 or 60 minutes using a motorized stage. For single cell traction experiments, cells were imaged using a 20X objective (NA 0.75 DIC, air) for a single time point. Traction forces were computed using Fourier-transform traction microscopy with finite gel thickness from a gel displacements field as previously described[33]. Gel displacements were obtained using a custom-made particle image velocimetry (PIV). In brief, the fluorescent beads in any experimental timepoint were compared to a reference image obtained after cell trypsinization at the end of the experiment.

**Kymographs and averaging**

To perform radial averages of traction forces, cancer cell clusters were segmented at every timepoint (either manually or automatically if cancer cells were fluorescently labeled). We then calculated the shortest (signed) distance of each pixel of the image to the cluster edge. Furthermore, we calculated the normal direction respect to the cluster edge as described previously[33] to decompose tractions in radial and tangential components. For every timepoint, we averaged each of these components according to their distance to the cluster edge to build spatiotemporal kymographs.

Different cancer cell clusters exhibit slightly different sizes due to experimental variability, hindering the averaging of traction profiles. To avoid averaging artifacts, each traction profile was linearly resized to the average cluster radius of the experimental condition before averaging.

**Pillar microfabrication**

The experimental pipeline to generate PAA microfabricated pillars is summarized in the Extended figure 7. At first, a negative SU8-50 photoresist (MicroChem) is coated onto a silicon wafer surface and patterned by conventional photolithography. In particular, an array of circular wells (50 μm radius, 50 μm deep) was fabricated using a chromium photomask (produced by direct laser lithography, Heidelberg μPG101) and the exposure-masking system UV-KUB2 (Kloé), following the photoresist manufacturer protocol. The well depth and surface roughness were verified by optical profilometer (Veeco). The SU8 master mold was then separated into individual array sub-units using a diamond scribe. To coat the flat surface of the SU8 master, 1 $cm^2$ PDMS stamps (~3 mm thick) were washed with 70% ethanol, rinsed with distilled water, and incubated for 1 h with 200 μl of a 100 μg/mL fibronectin solution (Corning). The excess of fibronectin was then aspirated and the PDMS surface was dried out thoroughly. SU8 masters were plasma treated for 1 min at high power (Harrick Plasma). Fibronectin-coated PDMS stamps were then put in contact with the SU8 plasma-treated surface and pressed down for 30 secs to allow fibronectin transfer between surfaces. Effective transfer was verified by checking the presence of circular non-transferred areas in the PDMS stamp in an inverted phase contrast microscope.

Once coated with fibronectin, SU8 masters were used to microfabricate PAA pillars. For this, 33 mm bottom-glass dishes (World Precision Instruments) were treated as before (Silane + Glutaraldehyde). PAA gels of 11kPa (Young modulus) were produced using a solution of 7.5% (v/v) acrylamide, 0.1% (v/v) bis-acrylamide, 0,5% (w/v) ammonium persulphate, 0,05% (w/v) tertamethylethylenediamine, 0,6% (w/v) Acrylic acic N-hydroxysuccinimide ester (stock 10 mg/mL in DMSO) and 2% (v/v) of 200-nm-diameter red fluorescent carboxylate-modified beads, in PBS. A 25 μl drop of this solution was added to the treated glass and the SU8 master was placed on. PAA gels were allowed to polymerize for 1h at RT. After, 3 ml of PBS were added and incubated for 30 minutes to facilitate lifting-off of the SU8 master, which was achieved with the help of a scalpel. Gels were then washed once with PBS and incubated with a solution of 100 μg/mL of fibronectin for 1h at RT to improve cell attachment. Gels were then washed twice with PBS and used immediately or stored at 4°C for a maximum of 24 h.

**Pillar compression assay**

To assess CAF pillar compression, a 100 μl drop containing ~200.000 pre-stained CAFs (CellTracker Green, ThermoFisher Scientific) was added on top of each pillar-containing PAA gel and incubated for 30-60 mins at 37°C, 5% $CO_2$ to allow attachment of cells. Non-attached cells were rinsed with culture medium and 3 mL of culture medium supplemented with 10%

(v/v) FBS, 1% (v/v) Insulin-Transferrin-Selenium (ThermoFisher Scientific), 2% (v/v) Antibiotic-Antimycotic (Gibco), and Metronidazol/Ciprofloxacin (20 and 5 μg/mL, respectively). Cells were then incubated overnight at 37°C, 5% $CO_2$.

Pillars were imaged on a spinning disk microscope (Nikon Eclipse Ti-E) equipped with thermal, $CO_2$, and humidity control, using a 20X objective (NA 0.75 Water) and a Z-step of 0,5 μm. A reference stack was acquired for each position after cell trypsinization at the end of the experiment. In drug perturbation experiments, pillars were initially imaged and then drugs were added on-stage (20 μM Para-Nitroblebbistatin (Optopharma), 50 μM Y27632 (SigmaAldrich) and incubated for 2 h. The same pillars were then re-imaged and a final reference stack was obtained after cells trypsinization.

**Pillar compression analysis**

The CAF-induced deformation of each pillar surface was quantified using a custom-made 3D PIV in Matlab[13]. The contact area between CAFs and the pillar was determined as the intersection of their respective fluorescent signals. The 3D traction field applied on the pillar was calculated, from the 3D deformation field, through a Finite Element Method by using ABAQUS (Dassault Systemes), in the large deformation regime. We prescribed the 3D displacement field as the boundary condition for the nodes that are at the contact area between the CAFs and the pillar, and the rest of the boundary was considered traction free, effectively constraining the tractions to the contact region. The pillar was modeled as a Neo-Hookean hyperelastic material. The projection of the tractions in the normal and tangential directions, unwrapping of the traction maps on the surface of the pillar, post-processing and averaging were performed in Matlab.

#### Supplementary References


25. Chanrion, M. *et al.* Concomitant Notch activation and p53 deletion trigger epithelial-to-mesenchymal transition and metastasis in mouse gut. *Nat. Commun.* **5**, 5005 (2014).
26. Jacobelli, J. *et al.* Confinement-optimized three-dimensional T cell amoeboid motility is modulated via myosin IIA-regulated adhesions. *Nat. Immunol.* **11**, 953–961 (2010).
27. Wendling, O., Bornert, J. M., Chambon, P. & Metzger, D. Efficient temporally-controlled targeted mutagenesis in smooth muscle cells of the adult mouse. *Genesis* **47**, 14–18 (2009).
28. Muzumdar, M. D., Tasic, B., Miyamichi, K., Li, L. & Luo, L. A global double-fluorescent Cre reporter mouse. *Genesis* **45**, 593–605 (2007).
29. Staneva, R., Barbazan, J., Simon, A., Vignjevic, D. M. & Krndija, D. Cell Migration in Tissues : Explant Culture and Live Imaging. in *Methods in Molecular Biology* **1749**, 163–173 (2018).
30. Calvo, F., Hooper, S. & Sahai, E. Isolation and Immortalization of Fibroblasts from Different Tumoral Stages. *Bio-protocol* **4**, e1097 (2014).
31. Zhao, J. J. *et al.* Human mammary epithelial cell transformation through the activation of phosphatidylinositol 3-kinase. *Cancer Cell* **3**, 483–495 (2003).
32. Pérez-González, C. *et al.* Active wetting of epithelial tissues. *Nat. Phys.* **15**, 79–88 (2019).
33. Brugués, A. *et al.* Forces driving epithelial wound healing. *Nat. Phys.* **10**, 683–690 (2014).

**Supplementary Table 1.**

| Antigen | Antibody/Chemical | Dilution | | | | | Reference | Vendor |
|---|---|---|---|---|---|---|---|---|
| | | FACS | cryosections | whole-mount | cultured cells | western blot | | |
| Fibronectin | Anti-Fibronectin rabbit polyclonal antibody | - | 1 to 100 | 1 to 100 | 1 to 200 | 1 to 10000 | F3648 | Sigma Aldrich |
| N-Cadherin | Anti-N-Cadherin mouse monoclonal antibody | - | - | - | - | 1 to 500 | 33-3900 | Thermo Fisher Scientific |
| MyosinIIA | Anti-MyosinIIA rabbit polyclonal antibody | - | - | 1 to 100 | 1 to 100 | 1 to 100 | 909801 | Biolegend |
| GAPDH | Anti-GAPDH rabbit polyclonal antibody | - | - | - | - | 1 to 5000 | G9545 | Sigma Aldrich |
| αSMA | Anti-αSMA mouse monoclonal antibody | 1 to 200 | - | - | 1 to 100 | - | A2547 | Sigma Aldrich |
| pMLC | Anti-pMLC rabbit polyclonal antibody | - | - | 1 to 100 | 1 to 200 | - | 3674 | Cell Signaling |
| YAP | Anti-YAP rabbit monoclonal antibody | - | 1 to 200 | - | 1 to 200 | - | 14074 | Cell Signaling |
| EpCAM | PE/Cy7 anti-human CD326, mouse IgG2b, κ | 1 to 10 | - | - | - | - | 324221 | Biolegend |
| EpCAM | APC anti-mouse CD326, mouse, rat IgG1 | 1 to 10 | - | - | - | - | 130-102-969 | Miltenyi Biotec |
| CD45 | Brilliant-Violet 421 anti-human CD45, mouse IgG1, κ | 1 to 20 | - | - | - | - | 304032 | Biolegend |
| CD45 | Brilliant-Violet 421 anti-mouse CD45, rat IgG2b, κ | 1 to 20 | - | - | - | - | 103133 | Biolegend |
| CD31 | FITC anti-human CD31, mouse IgG1, κ | 1 to 20 | - | - | - | - | 303104 | Biolegend |
| CD31 | FITC anti-human CD31, rat IgG2a, κ | 1 to 20 | - | - | - | - | 102405 | Biolegend |
| Vimentin | APC anti-human Vimentin, recombinant IgG1 | 1 to 250 | - | - | - | - | 130-106-370 | Miltenyi Biotec |
| Isotype controls | FITC mouse IgG1 | 1 to 10 | - | - | - | - | 130-092-213 | Miltenyi Biotec |
| | Brilliant-Violet 421 rat IgG2b, κ | 1 to 20 | - | - | - | - | 400639 | Biolegend |
| | FITC rat IgG2a, κ | 1 to 20 | - | - | - | - | 400505 | Biolegend |
| Anti-mouse Alexa Fluor 488 | Goat anti-mouse IgG Alexa Fluor 488 polyclonal antibody | - | 1 to 200 | 1 to 200 | 1 to 400 | - | A11029 | Thermo Fisher Scientific |

| | | | | | | | | |
|---|---|---|---|---|---|---|---|---|
| Anti-mouse Alexa Fluor 546 | Goat anti-mouse IgG Alexa Fluor 546 polyclonal antibody | - | 1 to 200 | 1 to 200 | 1 to 400 | - | A11030 | Thermo Fisher Scientific |
| Anti-Rabbit Alexa Fluor 488 | Goat anti-Rabbit IgG Alexa Fluor 488 polyclonal antibody | - | 1 to 200 | 1 to 200 | 1 to 400 | - | A32731 | Thermo Fisher Scientific |
| Anti-Rabbit Alexa Fluor 568 | Goat anti-Rabbit IgG Alexa Fluor 568 polyclonal antibody | - | 1 to 200 | 1 to 200 | 1 to 400 | - | A11011 | Thermo Fisher Scientific |
| Anti-rabbit HRP | Goat anti-Rabbit polyclonal antibody | | | | | | 32260 | Thermo Fisher Scientific |
| | Against Fibronectin | - | - | - | - | 1 to 10000 | | |
| | Against MyosinIIA | - | - | - | - | 1 to 5000 | | |
| | Against GAPDH | - | - | - | - | 1 to 2500 | | |
| Anti-mouse HRP | Goat anti-Mouse polyclonal antibody | | | | | | 32230 | Thermo Fisher Scientific |
| | Against N-Cadherin | - | - | - | - | 1 to 500 | | |
| F-actin | Phalloidin-Rhodamine | - | 1 to 200 | 1 to 100 | 1 to 200 | - | R415 | Thermo Fisher Scientific |
| | Phalloidin-Alexa Fluor 488 | - | 1 to 200 | 1 to 100 | 1 to 200 | - | A12379 | Thermo Fisher Scientific |
| | Phalloidin-Alexa Fluor 633 | - | - | - | 1 to 50 | | A22284 | Thermo Fisher Scientific |
| DNA | DAPI (4',6-Diamidino-2-Phenylindole, Dihydrochloride) | - | 1 to 400 | 1 to 400 | 1 to 400 | - | D1306 | Thermo Fisher Scientific |

**Supplementary Video 1. CAFs form a 3D capsule around cancer cells in tumors.**

3D image of a N/p53 tumor slice showing the organization of cancer cells (mGFP, green) and stroma (mTomato, magenta). The stroma forms intratumoral capsules that surround cancer cells and compartmentalize the tumor in clusters, as shown in the orthogonal views. Cell nuclei are labeled by DAPI (blue).

**Supplementary Video 2. CAFs compress cancer cells in vitro.**

<u>CAFs compression in cocultures</u>: Top (top) and lateral (bottom) views of the in vitro co-culture system showing a cluster of cancer cells (isolated from PDX, green) surrounded by CAFs (magenta). CAFs form a ring that closes on top of the cancer cells, reshaping the cluster and inducing the formation of a three-dimensional bud. White line indicates contour of the CAFs ring. Yellow line labels the location of the orthogonal view. Scale bar, 100 μm. Time, hh:mm.

<u>Imaris rendering of a 48h coculture</u>: 3D rendering of the CAFs-cancer cell coculture when the bud is stabilized (t=48h). Note that the CAFs ring is stalled around the bud, compressing cancer cells.

**Supplementary Video 3. Coculture of mouse cancer cells and CAFs.**

Phase contrast (left) and fluorescence of N/p53 cancer cells (right, mGFP) showing the evolution of the coculture. CAFs surround, reshape and compress cancer cells. Scale bar, 100 μm. Time, hh:mm.

**Supplementary Video 4. Spontaneous organization of CAFs and cancer cells from PDX tissue fragments**

Timelapse (phase contrast) of a coculture of cancer cells and CAFs that spontaneously exited a PDX tissue fragment. CAFs surrounded and confined cancer cells (right cluster). Some CAFs assemble a supracellular ring that reshapes and compresses cancer cells (bottom left cluster, yellow line). Scale bar, 100 μm. Time, hh:mm.

**Supplementary Video 5. Traction forces in cocultures**

Evolution of traction forces during the reshaping of the cancer cell cluster by CAFs. Left, DIC movie of the coculture outlining the contour of the cancer cell cluster (solid red line) and the

CAFs ring (dashed red line). Right, evolution of traction forces exerted by CAFs and cancer cells (black line labels the cancer cell cluster boundary). Tractions accumulate at the interface between cancer cells cluster and CAFs. At first timepoints, tractions point away from the cluster. When the CAFs ring is formed (t=9:00h) and during cluster reshaping, tractions point towards the cluster. Scale bar, 100 μm. Time, hh:mm.

**Supplementary Video 6. Inhibition of contractility suppresses compression of cancer cells in vitro**

Left, evolution of traction forces after treatment with blebbistatin. Right, evolution of traction forces after treatment with Y27632. Treatment starts at t=0h. Solid red line indicates the contour of the ring, dashed red line indicates the position of the ring immediately before treatment. Both treatments induce a massive decrease in traction forces and a fast relaxation of the CAFs ring. Scale bar, 100 μm. Time, hh:mm.

**Supplementary Video 7. Tissue displacements after laser ablation in tumors and in vitro cocultures.**

Left: laser ablation (t=0s) of the boundary between cancer cells (green) and the stroma (magenta) in a tumor slice. Ablated area is indicated in white. Tissue displacements are represented by white vectors. After ablation, cancer cells move towards the cut, showing that they are compressed. Solid cyan lines indicate the ROIs far from cut and near cut quantified in figure 2B. The cancer cell cluster is outlined by a dashed cyan line. Scale vector: 1 μm.

Center: laser ablation (t=0s) of the stroma (magenta) perpendicular to the boundary with cancer cells (green) in a tumor slice. Ablated area is indicated in white. Tissue displacements are represented by white vectors. After ablation, cancer cells move towards the cut, showing they are compressed, and the stroma recoils away from the cut, showing it is under tension. Solid cyan lines outline the ROIs of the stroma and cancer cells quantified in figure 2D. Yellow vectors show the average displacement in each ROI (for visualization purposes they are not scaled). The cancer cell cluster is outlined by a dashed cyan line. Scale vector: 1 μm.

Right: laser ablation (t=0s) of CAFs (magenta) perpendicular to the boundary with cancer cells (green) in vitro. Ablated area is indicated in white. Tissue displacements are represented by white vectors. After ablation, cancer cells move towards the cut, showing they compressed, and

CAFs recoil away from the cut, showing they are under tension. Solid cyan lines outline the ROIs of the CAFs and cancer cells quantified in Fig. 2F. Yellow vectors show the average displacement in each ROI (for visualization purposes they are not scaled). The cancer cell cluster is outlined by a dashed cyan line. Scale vector: 5 μm. Scale bar, 100 μm. Total time, 50 seconds.

**Supplementary Video 8. Tissue displacements after laser ablation in tumors with impaired CAFs contractility**

Laser ablations (t=0s) of the stroma perpendicular to the boundary with cancer cells (nGFP, green) in a control tumor (left), a blebbistatin treated tumor (center) and a tumor containing Myosin IIA KO CAFs (right). The stroma is labeled in magenta (mTomato) with the exception of Myosin IIA KO CAFs, which became green (mGFP) after myosin depletion. Ablated area is indicated in white. Tissue displacements are represented by white vectors. In the control tumor, cancer cells move towards the cut, showing they are compressed, and the stroma recoils away from the cut, showing it is under tension. This recoil pattern is lost when contractility is inhibited either by blebbistatin or by knocking-out myosin IIA in CAFs. Solid cyan lines outline the ROIs of the stroma and cancer cells quantified in figure 2G. Yellow vectors show the average displacement in each ROI (for visualization purposes they are not scaled). Cancer cell clusters are outlined by a dashed cyan line. Scale vector: 1 μm. Total time, 50 seconds.

**Supplementary Video 9. 3D visualization of a polyacrylamide pillar and CAFs organization**

<u>3D pillar visualization:</u> 3D rendering of fluorescent beads embedded in the polyacrylamide gel to visualize pillar 3D shape.

<u>3D visualization of CAFs in pillars:</u> CAFs organization around the pillar, visualized by immunostaining of F-actin (magenta), phospho-Myosin Light Chain (green) and nuclei (DAPI). CAFs form supracellular stress fibers rich in active myosin and align parallel to the pillar surface to compress it.

**Supplementary Video 10. CAFs traction forces exerted on the pillar**

Top (left) and side (right) view of a 3D pillar model in a relaxed state (frame 1), and progressively showing the deformations (orange) induced by CAFs traction forces (black vectors). Note that deformation is magnified 5 times for visualization purposes.

**Supplementary Video 11. Traction forces in cocultures of control and fibronectin-depleted CAFs**

Time-lapse of cocultures with control (top) and fibronectin-depleted (bottom) CAFs. Left, DIC image of the coculture outlining the contour of the cancer cell cluster (solid red line) and the CAFs ring (dashed red line). Right, evolution of traction forces exerted by CAFs and cancer cells (black line labels the cancer cell cluster boundary). Depletion of fibronectin impairs CAFs traction forces, ring formation and compression of the cancer cell cluster. Scale bar, 100 μm. Time, hh:mm.

#### Supplementary Note 1

In this note, we develop a physical model of the remodeling of the cancer cell cluster (CC cluster) by Cancer Associated Fibroblasts (CAFs) in the *in vitro* co-culture system. We aimed at providing a unified framework to quantitatively described the closure kinetics and the associated traction forces measured on the substrate, and to relate these to the pillar compression experiments.

### S1 Model

A sketch of the model system is presented in **Supp. Fig. 1a**. The CC cluster constitutes a cell monolayer prior to CAF remodeling. The closure of the CAF monolayer on top of the cluster is assumed to be solely driven by the contraction of the acto-myosin ring present at the inner edge of the CAFs, modeled as a constant line tension $\Gamma$ [1]. We concentrate on the dynamics characterized by inward traction forces, for which CAF migration may be disregarded. We further assume radial symmetry, which, although not particularly accurate in experimental situations such as the one shown in **Fig. 1d** of the main text, is a prerequisite for an analytically tractable model.

We adopt a thin sheet approximation for the mechanical description of the plane stress tensor $\gamma_{ij}$ in the CAF monolayer, and the stress tensor in the CC cluster, for which we only consider the integrated version over cluster thickness $h\sigma_{ij} \equiv \int_0^h \mathrm{d}z\, \sigma_{ij}$.

Several spatial regions must be distinguished, characterised by the radius of the CAF inner edge (the acto-myosin ring) $R_c$ and the outer edge of the CC cluster $R_T$ (see sketch on **Supp. Fig. 1a**).

- For $r > R_T$, the CAFs are in direct contact with the substrate and experience a substrate force density $T_r$. The local force balance in this region reads [2]:

$$\partial_r \gamma_{rr}(r) + \frac{\gamma_{rr}(r) - \gamma_{\theta\theta}(r)}{r} = T_r(r). \tag{S1}$$

  where $\gamma_{rr}$ and $\gamma_{\theta\theta}$ are the radial and orthoradial components of the CAF stress tensor and $r$ is the radial coordinate.

- For $R_c + \delta < r < R_T$, where $\delta$ is the width of the actomyosin ring, the CAFs are in contact with the CC cluster. The sliding of the CAFs on top of the CC cluster creates a shear stress $f(r)$ which acts as a source of tension gradient in both CAFs and CC cluster:

$$\partial_r \gamma_{rr}(r) + \frac{\gamma_{rr}(r) - \gamma_{\theta\theta}(r)}{r} = f(r)\ ,\ \partial_r h\sigma_{rr}(r) + \frac{h\sigma_{rr}(r) - h\sigma_{\theta\theta}(r)}{r} = -f(r) + T_r(r) \tag{S2}$$

- For $R_c < r < R_c + \delta$, the balance between the inward force from the ring line tension $\Gamma/R_c$ and the CAF radial tension may involve a particular lineic friction force $f_\Gamma$, which in turn causes a discontinuity $\delta h\sigma_{rr}$ in the CC cluster stress :

$$\gamma_{rr}(R_c) - \frac{\Gamma}{R_c} = f_\Gamma\ ,\ \delta h\sigma_{rr} = -f_\Gamma + T_r(R_c)\delta \tag{S3}$$

- For $r < R_c$, the central part of the CC cluster is not covered by the CAFs. Prior to bud formation, the central cluster is weakly deformed and does not directly exert stress on the actomyosin ring. The force balance in the CC cluster simply reads

$$\partial_r h\sigma_{rr}(r) + \frac{h\sigma_{rr}(r) - h\sigma_{\theta\theta}(r)}{r} = T_r(r) \tag{S4}$$

The persistence of inward pointing traction forces after the ring has reached a stationary position (**Fig. 1e**, main text, reproduced in **Supp. Fig. 1b**) suggests an elastic description for the mechanics of CAFs and CC cluster, and an elastic-like interaction with the substrate[1, 2]. For a 3D-incompressible elastic description of the CAF monolayer characterised by a 2D elastic modulus $E_c$, the components of the stress tensor are related to the radial displacement $u(r)$ in the CAF layer (where $r$ is the radial coordinate) according to:

$$\gamma_{rr}(r) = \gamma_c + \frac{2}{3}E_c\left[2\partial_r u(r) + \frac{u(r)}{r}\right] \;,\; \gamma_{\theta\theta}(r) = \gamma_c + \frac{2}{3}E_c\left[2\frac{u(r)}{r} + \partial_r u(r)\right] \tag{S5}$$

where $\gamma_c$ is the homeostatic contractile tension in the CAF monolayer far from the ring. Further assuming an elastic interaction with the substrate: $T_r(r) = Y_s u(r)$ where $Y_s$ is a static friction coefficient, the solution of Eq. 1 for the CAF displacement in the region $r > R_T$ is $u(r) \propto K_1(r/\lambda_s)$ where $K_1$ is a modified Bessel function, which leads to a spatial localisation of the substrate stress over a characteristic length scale $\lambda_s \equiv \sqrt{4E_c/(3Y_s)}$ [2]. This analytic solution is likely to be of limited validity in the present context, as the tissue mechanics is more complex than a simple elastic sheet, and include cellular rearrangements, especially during bud formation. However, the localisation of the traction forces outside the CC cluster is indeed observed (with $\lambda_s \simeq 50\ \mu$m), which suggests that the spatial derivative of the stress dominate the RHS of Eqs. 1,2,4 (quasi 1D approximation). These three local force balances can be integrated and added to Eq. 3 to obtain the global force balance

$$\frac{\Gamma}{R_c} = \gamma_c - \int_0^\infty T_r(r)dr - \gamma_{\text{bud}} \qquad \gamma_{\text{bud}} = h\sigma_{rr}|_{r=0} \tag{S6}$$

The LHS of this equation represents the force (per unit length) driving CAF closure and the RHS the forces resisting closure, which includes the CAF homeostatic tension, substrate friction, and the compression of the central region of the CC cluster. The expression given for $\gamma_{\text{bud}}$ corresponds to the low deformation limit of Eqs.1-4. However, Eq. 6 remains valid when a tumor bud forms in the central region tof the CC cluster, provided the bud compressive stress $-\gamma_{\text{bud}}$ is calculated by taking into account cellular rearrangement in the bud (see below).

#### S2 Traction force and closure dynamics

Eq. 6 may be seen as a dynamic equation for the evolution of the ring radius $R_c(t)$ as all the terms present in it (apart from $\gamma_c$) depend on time. The LHS of Eq. 6 clearly increases with time as the radius shrinks, so must the RHS. We may assume that the CAF tension far from the cluster $\gamma_c$ remains constant over time, so the increase of the closure force must be compensated by either an increase of traction force or by an increase of the bud resistance to compression or by both. **Supp. Fig. 1c,d** shows the temporal evolution of the total traction force [1] and the average ring radius (obtained from the ring area shown **Fig. 1f** in the main text). Two dynamical regimes can be distinguished from the moment where the total traction force points inward (**Supp. Fig. 1d**). At earlier time (from 7 to 16 hrs), the closure dynamics is fast ($dR_c/dt = -3.3\ \mu$m/h) and corresponds to a linear increase of the inward traction force with time. At later time (16 to 27 hrs), the closure dynamics is slower ($dR_c/dt = -1\ \mu$m/h) and the traction force saturates to a value $R_c \int_0^\infty T_r dr = 1.6\mu$N.

The linear increase of inward traction force at short time is consistent with the increase of the driving force from the line tension $\Gamma/R_c$ as the ring closes, although part of the increase of inward traction force must probably also be attributed to the decrease of active outward

[1] Two alternative definitions: $\int \mathrm{d}r\, rT_r$ and $R_c \int \mathrm{d}r\, T_r$, give similar results (**Supp. Fig. 1c**) which supports the approximation of localised tractions made to obtain Eq. 6.

traction force associated to CAF migration. The fact that the traction force saturates while the ring is still closing is remarkable, and indicates a build-up of stress in the central cluster region that is eventually driving bud formation. In this regime, only a fraction of the driving force is compensated by the traction force, which, at their saturation value, amounts to an effective line tension $\Gamma_s \simeq 1.2 - 1.6\mu$N. The remaining driving force must be compensated by the compression of the CC cluster. Thus, the CC cluster budding has a signature both in the closure dynamics and in the evolution of the traction force.

We can attempt to relate this value of the line tension to the stress measured in the pillar compression experiment reported in **Fig. 4** - main text. We see from **Fig. 4f** that the normal stress on pillars of radius $R_p = 50$ µm reaches $\sigma_p = 10^3$ Pa. This corresponds to a typical lateral tension in the CAF layer surrounding the pillar $\gamma_{\rm p} = \sigma_{\rm p} {\rm R}_{\rm p} = 5 \times 10^{-2}$ N/m. Multiplying $\gamma_p$ by the typical thickness $h_c \simeq 10$ µm of the CAF monolayer yields an estimate for the line tension $\Gamma \simeq 500$ nN. This is slightly smaller but of the same order of magnitude as the estimate obtained from the traction force in the co-culture experiments. This shows that the CAFs are indeed able to exert a large compressive stress, when compared, for instance, to the typical tension of one stress fiber, which is of order 10 nN [3].

#### S3 Bud stability

The cancer cell buds remain stable over time (**Fig. 1e,f** - main text and **Supp. Fig. 1b**) when directly compressed by the CAF ring. This experimental observation is not straightforward from a theoretical perspective. Indeed, the cellular rearrangements and multilayering involved in bud formation suggests a stress-dependent fluidification of the CC cluster. If the cluster behaved as a fluid, the bud would adopt a spherical shape due to its apical tension $\gamma_{cc}$. Experimental observations show that buds are rather cylindrical with small height-to-diameter ratio (**Extended Data Fig. 4b,c**). Furthermore, the stability of such fluid bud pinched by the actomyosin ring would rely on the ratio of driving to resisting force $\Gamma/(\gamma_{cc}R_c)$ to be of order unity. With a typical CC apical tension $\gamma_{cc} \sim 10^{-3}$ N/m and with $R_c \simeq 100$ µm, this ratio is about 10 times smaller.

To simplify the discussion of bud stability, we neglect the traction forces which saturate to a constant value after bud formation. We describe the bud as a stratified structure composed of $n$ layers (**Supp. Fig. 1e**), and we only discuss the diagonal component of the stress tensor in each layer, defined as a pressure $P$. To account for the plastic cellular rearrangements within the bud, we introduce a yield pressure $P^*$ that describes phenomenologically the amount of compressive stress necessary to trigger cell exchange between layers. CAF closure initially occurs on a monolayer of CCs, and force transmission through friction ($f(r)$ and $f_\Gamma$ in Eqs.2,3) leads to the generation of a compressive stress $P_1$ in the central part of the CC monolayer. If the pressure remains below the threshold ($P_1 < P^*$), no multilayering occurs and the CAFs are able to close on top of the CC monolayer, as it is sometimes observed in our experiments (data not shown). If the threshold is exceeded, cells delaminate from the central region of the CC cluster and form a second layer. The particular ring radius at which this happens is defined as $R_c^*$. The pressure $P_2(t)$ in the second layer increases as the ring closes and cells are transferred from the first layer. If $P_2 > P^*$, a third layer forms, with pressure $P_3(t)$, and cells are expelled from the second payer until $P_2 - P_3 < P^*$. Only the second layer is directly compressed by the ring, so the bud resistive force in Eq. 6 is $\gamma_{\rm bud} = -hP_2$. The pressure $P_2$ is estimated as follows. For $P_2 < P^*$, $P_2 = E_{cc}(R_2^* - R_c)/R_2^*$, where $E_{cc}$ is the bulk modulus of the layer and $R_2^*$ is the ring radius at which layer 2 starts being compressed (**Supp. Fig. 1f**). For $P_2 > P^*$, $P_2 = P^*\phi(h_b/h)$, where $h_b$ is the height of the bud, determined by volume conservation ($h_b = h(R_2^*/R_c)^2$) and $\phi(x)$ is a phenomenological shape function capturing the shear stress between adjacent layers. In analogy with the standard problem of elastic pillar deformation induced by a collar pressure [4], we adopt the function $\phi(x) = 1 + b\tanh[(x-1)/b]$ which appropriately describes the localisation of the stress near the ring for tall bud ($h_b$), as seen in the pillar compression experiment (**Fig. 4g** - main text). Here, $b$ is a phenomenological parameter representing shear resistance between adjacent layers. For an homogeneous elastic material, finite element simulations yield $b \simeq 0.3$. In the present context, $b$ is determined by cell-cell adhesion and can be considered as a free parameter.

**Supp. Fig. 1f** shows a graphical representation of the static force balance between the ring tension and the bud compression $\Gamma/R_c = \gamma_c + hP_2$. The existence of a static equilibrium with a finite ring radius state crucially depends on the parameter $b$. If the ring tension is shared by all layers ($b \to \infty$), there is always a single stable equilibrium as the height of the bud diverges when $R_c$ vanishes (solid gray line in **Supp. Fig. 1f**). In the opposite limit ($b \to 0$) different layers are uncoupled, and $P_2 \leq P^*$. Since the formation of a second layer requires $\Gamma/R_c > \gamma_c + hP^*$, no static equilibrium exists at finite $R_c$ and multilayering necessarily leads to ring closure below the bud (dashed gray line in **Supp. Fig. 1f**). Under more realistic situations, bud compression leads to a maximum resistive force $hP^*(1+b)$ that defines three phenotypes according to the parameter values (**Supp. Fig. 1e,f**): full budding and CAF closure, stable bud with a finite neck size and CAF closure with no bud. This is summarized on **Supp. Fig. 1g** with a phase diagram as a function of reduced yield pressure $P^*/E_{cc}$ and reduced line tension $\Gamma/(E_{cc}h^2)$. If $P^*$ is small, the CAF shear stress triggers the multilayering transition for large ring radius but the maximum bud resistive force is too small to equilibrate the ring compression. If $P^*$ is large, the multilayering transition occurs for a small ring radius, for which ring compression increases faster than the bud resistive force with decreasing ring radius. In both cases, cells are expelled from the second layer until the CAF ring closes. For even larger $P^*$, multilayering would only occur for a ring radius of order the cell size, in which case we consider that the ring closes without bud formation (gray region on **Supp. Fig. 1g**). Thus, bud stability occurs only on a parametric region were both ring line tension and yield pressure are sufficiently small, and can be enhanced with an increase of CC elastic modulus $E_{cc}$ or effective parameter $b$. From the estimation $\Gamma \sim 1$ μN obtained in the previous part and with $h \sim 10$ μm, stable buds require $E_{cc} \gtrsim 1$ kPa and $P^* \simeq 0.1 - 0.4E_{cc}$. Thus, CAF compression around an incomplete CC bud provides an indirect way to measure the tissue elastic modulus, found here to be consistent with typical values measured for cancer cells [5].

### References


[1] S. R. K. Vedula, G. Peyret, I. Cheddadi, T. Chen, A. Brugués, H. Hirata, H. Lopez-Menendez, Y. Toyama, L. Neves de Almeida, X. Trepat, C. T. Lim, and B. Ladoux, "Mechanics of epithelial closure over non-adherent environments," *Nature Communications*, vol. 6, no. 1, p. 6111, 2015.

[2] C. M. Edwards and U. S. Schwarz, "Force Localization in Contracting Cell Layers," *Phys. Rev. Lett.*, vol. 107, no. 12, p. 128101, 2011.

[3] S. Deguchi, T. Ohashi, and M. Sato, "Tensile properties of single stress fibers isolated from cultured vascular smooth muscle cells," *Journal of Biomechanics*, vol. 39, no. 14, pp. 2603–2610, 2006.

[4] V. V. Meleshko and Y. V. Tokovyy, "Equilibrium of an elastic finite cylinder under axisymmetric discontinuous normal loadings," *J Eng Math*, vol. 78, no. 1, pp. 143–166, 2013.

[5] C. Alibert, B. Goud, and J.-B. Manneville, "Are cancer cells really softer than normal cells?," *Biology of the Cell*, vol. 109, no. 5, pp. 167–189, 2017.

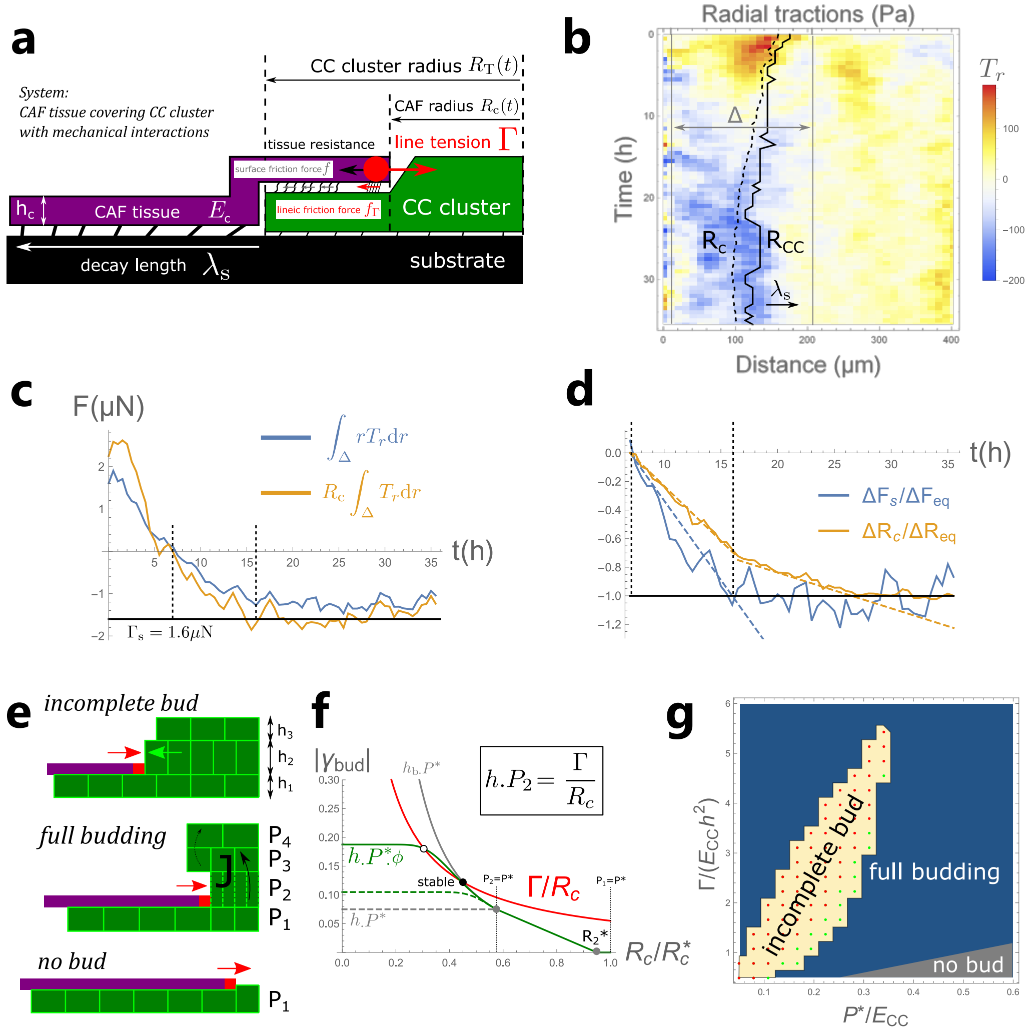


**Supplementary figure 1** **(a)**: Sketch of the model: a CAF monolayer (purple) closes on top of a cluster of cancer cells (green), driven by the line tension provided by a dense actomyosin ring (red). The CAFs exert a surface friction force on the CC cluster – which may include a lineic friction $f_\Gamma$ at the location of the ring – that creates cluster compression. The movement of both the CAFs and the CC cluster generate traction forces on the substrate. Within an elastic model, the traction force below the CAF layer are localised within a decay length $\lambda_s$. **(b)**: Experimental kymograph of substrate tractions $T_r$ (reproduced from **Fig. 1e** - main text) showing the localization on length $\lambda_s$. Dashed and solid black lines respectively represent the location of the actomyosin ring and CC cluster periphery. We only focus on times for which tractions are inward (blue). The ROI is defined as the spatial range $\Delta$ (gray lines). Outside this range, numerical ($r = 0$) or biological ($r > 200$ µm) fluctuations are dominant. **(c)**: The computed total traction force over the ROI, through a surface integral $\int_\Delta rT_r(r)\mathrm{d}r \equiv F_s/(2\pi)$ (blue) or a line integral $R_c\int_\Delta T_r(r)\mathrm{d}r$ (orange). Inward tractions (negative) start after $t = 7$ h and reach a saturation value between 1.2 and 1.6 µN for $t > 16$ h. **(d)**: Relative variation of the total traction force and the ring area from their value at $t = 7$ h, normalised by their saturation value at long time. A striking feature is the apparent saturation of traction force before CAF edge radius saturation, that we interpret as an indication of bud compression. For all studied clusters with stabilized buds, abrupt changes in ring and traction dynamics coincide. **(e)**: Sketch of the plastic model of bud formation: CC cluster multilayering occurs if the stress within the cluster generated by CAF closure exceeds a threshold $P^*$, and cell transfer between adjacent layers occurs if the stress difference between the two layers exceed $P^*$. The second CC layer directly resists the CAF ring tension. This may result in a stable incomplete bud with a finite neck size, or ring closure and full budding. **(f)**: Force balance between the CAF ring compression (red) and the bud elastic resistance (green), as a function of ring radius $R_c$. The shear-transmitted resistance from adjacent layers to second layer is captured by a shape factor $\phi = 1 + b\tanh[(h_b/h - 1)/b]$. A second layer is formed for a particular ring radius $R_c = R_c^*$ (when $P_1 = P^*$), and is compressed up to $P_2 = P^*$ and eventually builds more layers. This may lead to a stable incomplete bud (green) or to ring closure and full budding (dashed green) if bud resistance is too weak. The parameter $b$ characterises the shear stiffness between layers. The dashed and solid gray lines show the asymptotic cases $b \to 0$ and $b \to \infty$, respectively. **(g)**: Budding phase diagram as a function of the dimensionless yield pressure $P^*$ and ring line tension $\Gamma$ ($E_{cc}$ is the CC bulk modulus). A stable bud occurs only for a limited range of both parameters (yellow) and can contain two layers (green dots) or more than two (red dots). The "no bud" region (gray) corresponds to a bud that would contain less than two cells. Parameters used are $E_c = 0$, $b = 1.5$.
